# Do single-cell foundation models need 512 dimensions for one number?

**DOI:** 10.64898/2026.11.24.768376

**Authors:** Lecheng Zhang, Yuhao Yang, Stan Z. Li

## Abstract

Single-cell foundation models represent counts through high-dimensional value embeddings or ranked feature tokens. We asked how many dimensions counts need and what network computes. In a controlled Transformer, a four-dimensional value track separated from a wide gene-identity backbone retains nearly all masked-prediction net gain; a straight line replaces pretrained value curves with little transfer loss. Across evaluated cohorts, clamping detected-gene values retains most gain in annotation, surface-protein prediction and perturbation-shift consistency, consistent with the prevalence of single-molecule counts. A two-feature encoder combining tie-free rank and within-cell standardized magnitude improves controlled retraining and preserves annotation when grafted onto pretrained scGPT. Gene offsets and three linear cell-level scalars approximate masked prediction and predict responses to count thinning; depth leads in the controlled model, while cell type leads in scGPT. Matched controls show that attribution, attention, dictionary and probe readouts can reflect depth, gene frequency, hub degree or directions that transfer little output.

## Introduction

Single-cell foundation models are pretrained on tens of millions of cells to annotate cell types, integrate atlases and predict responses to perturbations [1–5]. Although each gene or peak contributes one scalar count, these Transformers represent each feature with hundreds of coordinates. Learned value encoders embed expression magnitudes in scGPT and scFoundation [1, 2], gene rank order conveys expression in Geneformer [3, 4], and soft-binned count embeddings are added to protein-derived gene features in State [5]. Shrinking an entire Transformer narrows the scalar value pathway and the gene-combining backbone together, leaving open how many dimensions the count itself needs and what the network computes from it.

Independent benchmarks have reported modest or task-dependent advantages of pretrained models over linear and task-specific baselines [6–9], with clearer gains emerging on calibrated metrics that account for technical noise [10–12]. Explaining when value representations help requires examining the measurements they encode. Droplet transcriptomes are sparse and contain many low positive counts, making feature detection an important component of their variation [13, 14], while scGPT’s within-cell quantile binning splits tied counts across bins at random. Recent studies have compared learned gene tables with protein-language-model embeddings [15], explored raw-count inputs [16] and tested how pretraining data scale and diversity affect performance [17], yet it remains unclear how much downstream tasks rely on expression magnitudes beyond detection, or whether a compact encoder built around tie-free ranks and relative magnitudes can replace wide nonlinear value networks.

Even when a model predicts held-out counts accurately, prediction error alone does not identify the computation behind its score. Distinct predictors can achieve similar scores through different computations; in causal inference, distinct structures can also be observationally equivalent [18–20]. Inside pretrained networks, interpretability studies have probed biological coherence [21], scored attention weights against regulatory databases [22] and decomposed activations into sparse dictionaries and manifolds [23, 24]. Sequencing depth and gene prevalence structure the input data, while hub degree can dominate regulatory benchmarks. In addition, linearly decoding a state from hidden layers does not ensure that predictions route through the decoded directions. Separating learned regulation from these statistical effects therefore requires functional interventions and matched baselines.

Here we trace scalar measurements from input tokens to predictions and internal readouts in pretrained transcriptomic models (scGPT, scFoundation, Geneformer-V2 and State), chromatin-accessibility models (EpiAgent and ChromFound) and controlled Transformers trained from scratch. Evaluations span human tissue transcriptomes, paired surface-protein measurements, pooled genetic perturbations and single-cell chromatin-accessibility cohorts. Separating value transport from a wide gene-identity backbone allows a narrow value track to preserve most masked-prediction gain, while a two-feature encoder based on tie-free rank and within-cell standardized magnitude improves controlled retraining and preserves annotation during pretrained adaptation with far fewer value-encoder parameters. Across annotation, surface-protein prediction, perturbation consistency and chromatin accessibility, transfer relies mainly on which features are detected. Beyond cohort means, gene offsets plus a readout of a few linear cell-level scalars led by sequencing depth or coarse cell type approximate masked prediction and predict responses to count thinning. Matched controls show that attribution, attention, dictionary and probing readouts can reflect embedding-path geometry, hub degree, gene frequency or decodable state directions that transfer little output when swapped.

## Results

### The expression value needs a few dimensions

Narrowing the whole network and narrowing only the value path give opposite answers (Fig. 1a). When every component of an scGPT-style Transformer shares one width, the net gain over cohort means falls as the width shrinks, to 30% of the 512-dimensional gain at four dimensions, and cell-type annotation from the cell embedding falls with it. The two-track Transformer separates the two roles: gene identity keeps a wide track that routes attention and forms the cell embedding, while the expression value travels through all six layers on its own narrow track. A four-dimensional value track with 2,080 parameters keeps 95% of the net gain, and annotation changes by less than half a macro-F1 point (Fig. 1c–e and Extended Data Fig. 3). Whole-network narrowing loses gain in every tissue, whereas the four-dimensional value track retains 95% of the gain with 9,850-fold fewer value-track parameters (Fig. 1h,i). Because the wide identity track remains, total model parameters fall from 35.5 million to 15.0 million, a 2.36-fold reduction. Annotation meets the pre-specified 1.5-point equivalence margin in the 40-donor evaluation; the 24-donor interval extends just beyond the lower margin.

**Figure 1.**
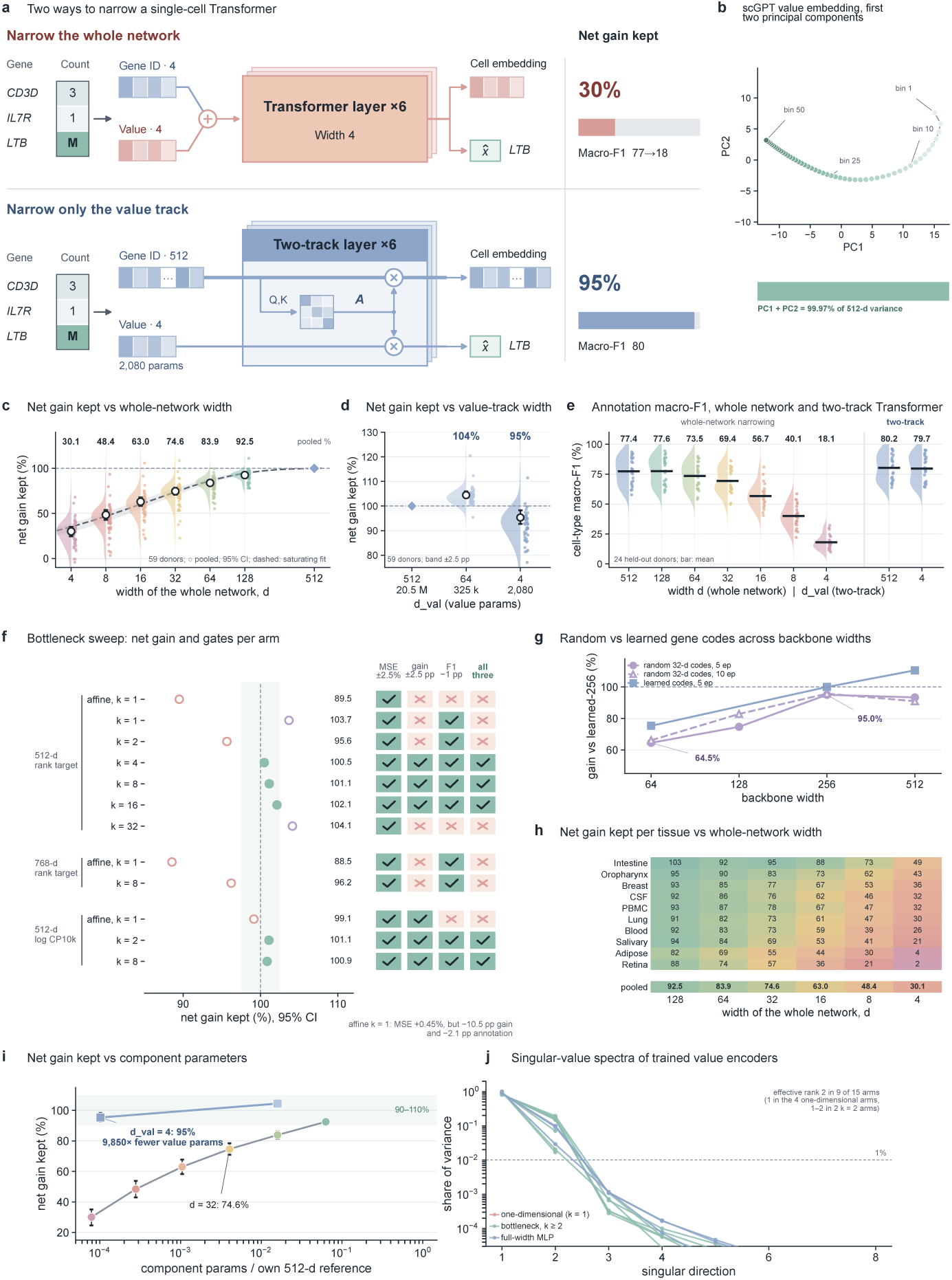
Value-track narrowing preserves net gain; whole-network narrowing reduces it. **a**, The schematic defines whole-network and value-track narrowing in controlled Transformers (Transformers trained from scratch under controlled conditions). Colours separate gene identity, values and masked queries. **b**, Pretrained scGPT value embeddings are projected onto principal components. Colour marks input bins; the bar partitions projected and remaining variance. **c**, Masked-prediction net gain is compared across network widths. Dots represent 59 held-out donors, averaged across three runs; rings and bars show pooled estimates and 95% donor-bootstrap intervals; the dashed fit has a 95% bootstrap band. **d**, Value-track narrowing is evaluated on the same donors. Rings show pooled estimates with 95% donor-bootstrap intervals; shading marks *±*2.5 percentage points around the wide reference. **e**, Cell-type annotation compares both architectures. Dots represent 24 held-out donors; horizontal bars mark means. **f**, Bottleneck models compare net gain and acceptance criteria across three runs per arm under rank-target and log counts per ten thousand (CP10k) objectives. Filled markers pass all gates; bars show 95% bootstrap intervals, and tiles test error within *±*2.5%, gain within *±*2.5 points and annotation no lower than *−*1 point (Methods). A failed two-sided gain gate can reflect improvement: nonlinear K1 exceeds the upper margin while passing masked-prediction and downstream criteria. **g**, Random and learned gene codes are compared across backbone widths. Symbols distinguish code types and training durations; gain is relative to learned codes at the reference width. **h**, Tissue cohorts compare whole-network narrowing. Heat-map colour and labels encode relative net gain; the bottom row pools donors. **i**, Relative net gain is plotted against the fraction of narrowed parameters. Circles denote whole networks and squares value tracks; bars show 95% donor-bootstrap intervals and shading marks the 90–110% reference band. The horizontal axis uses different component denominators: backbone/value parameters in the whole-network scan and value-stream parameters in the two-track scan, each normalised to its own width-512 reference. Total two-track parameters fall from 35.5 to 15.0 million at value width four. **j**, Value-network spectra compare encoder configurations. Curves show squared three-run mean singular values, normalised over the plotted directions; colour groups encoder types and the vertical axis is logarithmic.

Pretrained value encoders already use few directions. Two principal components carry more than 99.9% of the variance of scGPT’s learned value curve and 97% of scFoundation’s (Fig. 1b). Replacing either curve with a straight line, an affine value encoder, changes downstream transfer little. Masked prediction by scFoundation on K562 cells is the exception: the straight line retains only 44.76% of its masked-prediction gain. Chromatin-accessibility models show the same geometry where they encode counts as values: ChromFound’s value embedding has an effective rank of one, whereas EpiAgent encodes accessibility through peak ranks and has a high-dimensional rank embedding (Extended Data Fig. 2k).

Training from scratch gives the same answer. Across ten cohorts evaluated on held-out donors (Extended Data Fig. 1 and Methods), an affine value encoder matches an MLP value encoder on masked prediction, annotation and surface-protein prediction, and the match holds at one million training cells (Extended Data Fig. 2c,d). An MLP value encoder that is free to use many directions collapses to an effective rank of two after training (Fig. 1j), so the nonlinearity buys little that the network keeps.

A bottleneck sweep evaluated against pre-specified acceptance criteria identifies several sufficient widths. In an scGPT-sized rank-target Transformer, bottlenecks of 4, 8 and 16 dimensions meet every criterion on masked-prediction error, net gain and downstream probes, and the one-dimensional affine value encoder keeps 90% of the net gain (Fig. 1f). The one-dimensional nonlinear bottleneck retains 103.7% of net gain and passes the error and downstream criteria, but fails two-sided gain equivalence because its improvement exceeds the upper margin. Thus, passing widths identify sufficient settings rather than a minimum dimension (Methods).

The gene-combining backbone also needs width. With fixed random gene codes, a 256-dimensional Transformer recovers 95% of the masked-prediction gain of learned codes, whereas narrower backbones recover less and learned codes pull ahead only at the widest setting (Fig. 1g). These results suggest that backbone width matters more than learned gene codes at intermediate widths.

### Downstream tasks read which genes are detected

To compare tasks on one scale, we gave each pretrained model four versions of every cell: the original values, a clamped input with one constant value on every detected gene, values shuffled among the detected genes, and permuted gene identities as the lower reference (Fig. 2a). Retention is the gain of an altered input above the permuted reference, as a share of the gain of the original input (Methods). Probes were refitted after each intervention. Across four models and five tasks, the three models that read expression through detected genes keep most of their gain under clamping (Fig. 2l); scFoundation retains its supplied depth tokens in this primary comparison, with fixed-depth controls reported separately.

**Figure 2.**
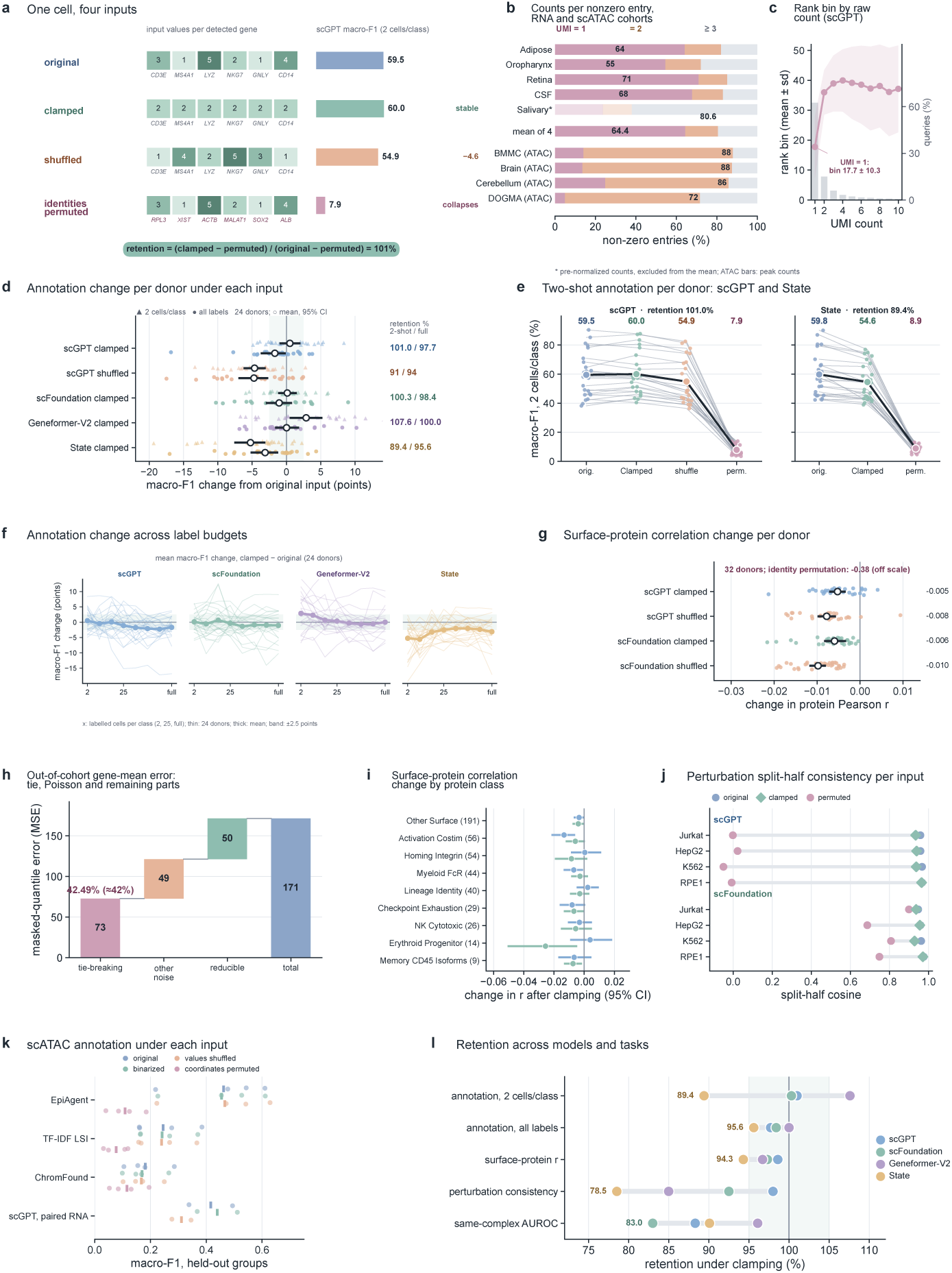
Gene detection carries most of the downstream gain. **a**, The schematic defines original input, clamped input, shuffled-value input and identity-permuted input. Colours distinguish interventions and gene labels identify detected transcripts. **b**, Count distributions compare tissue and chromatin-accessibility cohorts. Stacks separate one, two and higher counts; RNA counts are unique molecular identifiers (UMIs), peak counts are accessibility counts, and faded salivary counts are excluded from the RNA mean. **c**, Rank bins are grouped by UMI count. The line and band show mean *±* standard deviation; bars show query shares. **d**, Cell-type annotation compares interventions across 24 held-out donors. Triangles denote two labelled cells per class (two-shot), circles all labels, and rings means with 95% donor-bootstrap intervals. **e**, Pretrained scGPT and Arc State/State embedding model compare two-shot annotation. Thin lines connect donors; thick lines connect means. **f**, Clamping effects are compared across label budgets. Thin lines denote donors, thick lines means, and shading *±*2.5 points. **g**, Surface-protein prediction compares clamping and shuffling with cognate transcripts masked. Dots represent 32 donors; rings show means with 95% donor-bootstrap intervals. **h**, Gene-mean error is decomposed across five cohorts. Waterfall bars separate tie variance, additional Poisson-model noise and the remaining error (Methods). **i**, Protein classes compare clamping effects. Points show mean correlation changes with 95% protein-bootstrap intervals. **j**, Perturbation-response consistency compares input conditions. Symbols distinguish conditions within each cell line and model. **k**, Accessibility annotation compares EpiAgent, ChromFound, term frequency–inverse document frequency (TF-IDF) with latent semantic indexing (LSI), and paired-RNA scGPT. Dots mark cohorts; ticks mark means. Accessibility probes remain fixed after intact-input fitting; groups are donors in three cohorts and libraries in DOGMA. **l**, Retention compares models across tasks. Colour identifies models; shading marks 95–105% retention and complex recovery uses area under the receiver operating characteristic curve (AUROC).

Cell-type annotation keeps its gain without the values. Across 24 held-out donors from five cohorts, clamped inputs retain 98–101% of the gain in scGPT and scFoundation, from two labelled cells per class to full labels, and Geneformer-V2 retains at least as much. This average masks donor variation: scGPT changes range from -16.8 to +8.4 macro-F1 points, and full-label mean annotation falls by 1.7 points. Shuffling values costs scGPT about five macro-F1 points, showing that the original encoder uses magnitudes even though refitted probes recover most average transfer from clamped inputs. State, which encodes magnitudes more directly, serves as a positive control, and clamping lowers its two-shot retention to 89% (Fig. 2d–f).

Surface-protein prediction and the consistency of perturbation-associated embedding shifts show similar retention. When 463 proteins are predicted from expression with their cognate, paralogous and subunit transcripts masked, clamped inputs retain 97–99% of the gain in scGPT, scFoundation and Geneformer-V2, and the average effect remains small across protein classes (Fig. 2g,i). Across 320 CRISPRi perturbations in four cell lines, clamped inputs keep the split-half consistency of each perturbation-associated embedding shift, whereas permuting gene identities removes it (Fig. 2j). This measure is the split-half agreement of observed cell-subset shifts, not prediction on held-out perturbations. Recall of protein complexes keeps less of its gain, which shows that magnitudes contribute to complex structure, and partial fine-tuning does not consistently increase sensitivity to clamping. After fine-tuning, annotation changes by 0.61 and +1.23 macro-F1 points at full labels in scGPT and scFoundation, respectively; with ten labels per class, it falls by 1.79 and 1.94 points. No equivalence test was conducted for these contrasts (protein comparisons in Extended Data Fig. 4c).

Chromatin-accessibility models also read which features are present. In four scATAC-seq cohorts, binarizing peak counts or shuffling nonzero values within a cell changes EpiAgent annotation by at most about 4.3 macro-F1 percentage points across the evaluated pooling choices (three donor-grouped cohorts and a DOGMA library-grouped comparison with fixed intact-trained probes), whereas permuting peak coordinates removes about 70% of its score (Fig. 2k). TF-IDF with latent semantic indexing behaves the same way. Applied to the paired RNA of two of these cohorts, scGPT does lose annotation when its values are shuffled, so the same assay detects magnitude use where it exists.

The counts themselves explain why detection carries so much. In the four held-out tissue cohorts with raw counts, about two thirds of nonzero entries are single molecules, most nonzero scATAC entries are one or two, and detection alone accounts for 83% of the per-gene variance of log-normalized expression. Under scGPT’s within-cell quantile binning, genes with equal counts are split across bins at random, so a single molecule can land in almost any bin, and this tie-breaking accounts for 42% of the error of an out-of-cohort gene-mean predictor (Fig. 2b,c,h). In scFoundation, raw predictions improve on out-of-cohort gene means only when zeros are among the queries, and calibrating detection rates and visible depth recovers the gain on positive-only queries (Extended Data Fig. 4b).

Tie noise and depth motivate an encoder that represents tie-free rank and relative magnitude explicitly. The two-feature encoder combines a tie-free mid-rank with a within-cell standardized log magnitude in 1,280 parameters, against 66,816 for the MLP value encoder. Trained on ten cohorts and scored on 59 held-out donors, it lowers masked-prediction error against the quantile encoder in every donor and raises three-run mean annotation from 0.66 to 0.77 macro-F1 (0.68 to 0.79 for the three-run ensemble). Either tie-free feature alone captures most of the error reduction and annotation gain, whereas reintroducing raw depth into the classification token degrades both. Grafted onto pretrained scGPT, the two-feature encoder replaces 264,704 value parameters with 2,560 and keeps annotation at the level of the original encoder (Extended Data Fig. 2f,g); its advantage persists in a 12-layer Transformer trained on one million cells (Methods).

### A few cell-level scalars approximate much of masked prediction

Although downstream transfer reads mainly gene detection, masked prediction still uses visible context to predict held-out expression, and prediction accuracy alone does not establish how closely a surrogate approximates the Transformer’s outputs. For calibrated predictors with nested information, the share of the Transformer’s function a predictor reproduces, its distillation *R*^2^, equals the share of the gain it recovers. The controlled Transformers are calibrated in this sense, and predictors built from the sparse panel’s own statistics sit on this diagonal. Nearest-neighbour retrieval on the full vocabulary sits far below it: it recovers 81% of the gain while reproducing only 22% of the function, so a high score can come from a different computation (Fig. 3b). Agreement with the Transformer’s own outputs, and not only with the targets, is therefore the measure we use to describe what the Transformer computes.

**Figure 3.**
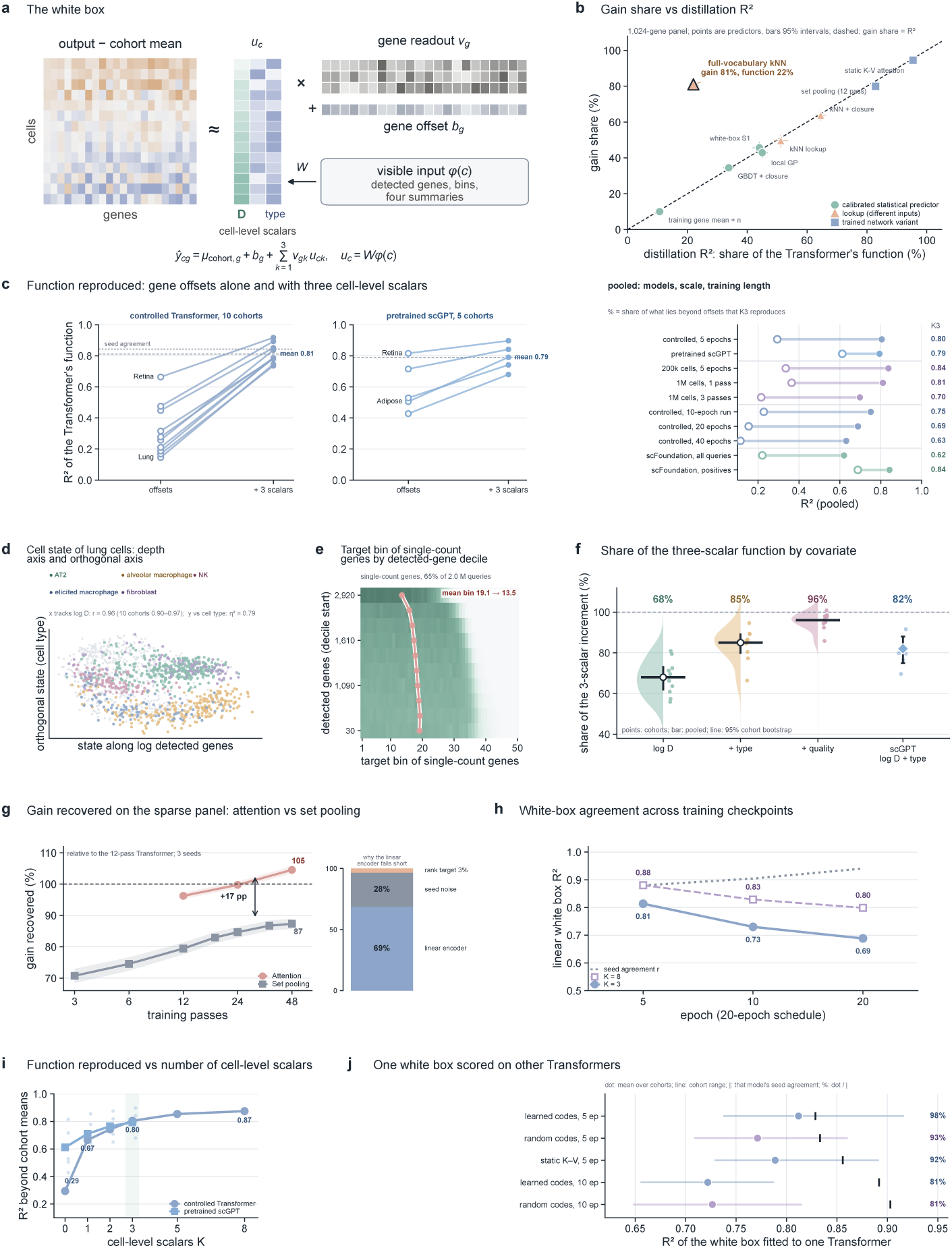
Gene offsets and a few cell-level scalars approximate much of masked prediction. **a**, The schematic defines a white box comprising cohort means, gene offsets and a readout of cell-level scalars. Coloured matrices distinguish cell state from gene readouts; the arrow denotes the visible-input map (Methods). **b**, Masked-prediction gain share is compared with distillation agreement. Symbols distinguish predictor families; bars show 95% run-by-cell bootstrap intervals and the diagonal marks equality. **c**, Offset-only and cell-state fits compare controlled Transformers with pretrained scGPT. Lines join each cohort’s fits, open and filled symbols distinguish them, dashed lines mark cohort means, dotted lines mark independent-model agreement, and right-hand dumbbells show pooled comparisons. **d**, Lung cells are displayed in depth-aligned cell-state coordinates. Colour marks cell type; the second axis is orthogonal to depth. **e**, Single-count target bins are grouped by detected-gene-count decile. Heat-map intensity shows frequency; the overlaid line marks mean bins. **f**, Depth, cell type and quality covariates are compared as predictors of cell state. Dots denote cohorts; summary markers show pooled shares with 95% cohort-bootstrap intervals. **g**, Sparse-panel masked prediction compares attention with set pooling over training. Bands show 95% cell-bootstrap intervals across three runs; the inset partitions the sparse-context shortfall among encoder, model noise and rank contributions. **h**, White-box agreement is compared across training checkpoints. Lines distinguish numbers of cell-level scalars; the dotted reference shows independent-model agreement. **i**, Function agreement is plotted against the number of cell-level scalars. Curves distinguish model families, small points mark scGPT cohorts, and shading highlights the reference dimension. **j**, A white box fitted to one Transformer is evaluated on other Transformers. Dots show cohort means, bars cohort ranges, and ticks each model’s independent-run agreement.

Beyond cohort means, we fit the following white-box approximation to the controlled Transformer (Fig. 3a):

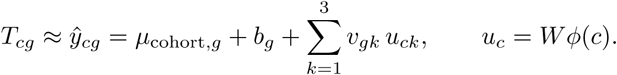

Each gene has an offset *b_g_* and a readout *v_g_* of a cell state *u_c_* with three cell-level scalars, and the cell state is a linear map of the visible input *ϕ*(*c*): which genes are detected, their bins and four summary statistics. On 59 held-out donors, the offsets alone reproduce 0.29 of the Transformer’s function beyond cohort means, and the three cell-level scalars raise this to 0.80, which is 95% of the agreement between independently trained runs (Fig. 3c,i). This ratio compares output agreement, not the fraction of an internal causal mechanism recovered. The same white box also describes Transformers trained with random gene codes, static key–value attention or longer schedules (Fig. 3j). In pretrained scGPT it reaches a similar agreement on five held-out cohorts, but the gene offsets carry more of the function, and the cell-level scalars describe 47% of what lies beyond them, against 76% in the controlled Transformer on the same cohorts.

The leading cell-level scalar is sequencing depth, measured by the detected-gene count *D*. In the controlled Transformer, log *D* alone accounts for 68% of what the three scalars add beyond the offsets, as much as total UMI, mitochondrial and ribosomal fractions combined, and coarse cell type raises the share to 85% (Fig. 3d,f). In pretrained scGPT, cell type and *D* together account for a similar share, with cell type leading. Depth therefore sets much of what the model predicts for every masked gene of a cell, and the model needs no explicit depth input to read it: although two thirds of its training cells arrive as a random window of 1,024 detected genes, the visible window predicts the whole-cell log *D* of held-out donors almost exactly (Extended Data Fig. 6b).

Under rank targets, the dependence on *D* follows from ranking itself. Each target ranks a gene among the genes its cell detected, so a gene’s expected bin moves with the number of genes that compete with it, and a parameter-free multinomial model that knows only average gene abundances and the binning rule predicts how strongly each gene responds to *D* (Fig. 3e and Extended Data Fig. 6c). Ranking targets within the input window instead of the whole cell leaves the share of *D* unchanged. No pathway enrichment was detected along the tested gene readouts: after controlling for abundance and detection rate, none of 1,801 Hallmark, Reactome and curated gene sets is enriched along them (Extended Data Fig. 6e).

Longer training adds cell-level scalars without replacing the leading ones. Five times more cells at the same number of optimization steps leaves the three-scalar description almost unchanged, with *D* still leading. Three passes over the larger corpus raise the net gain by 70%, and the function then needs more cell-level scalars: three reproduce 0.70 of it and eight reproduce 0.79, the same values a free per-cell state reaches (Fig. 3h and Extended Data Figs. 6d and 7a). Within the tested corpus sizes and schedules, additional optimisation steps increase the dimensionality needed by the approximation more than adding cells at fixed steps. This complements reports of rank growth in network weights and embeddings [25–28].

Pairwise attention earns its cost when context is sparse. On a panel of 1,024 genes with about 18 visible genes per cell, attention outperforms an architecture-matched set model by 17 percentage points of gain recovered. With so few visible genes the cell state stays low-dimensional, but a linear encoder no longer reaches it, and most of that shortfall comes from the encoder itself (Fig. 3g and Extended Data Figs. 6a and 7d).

### Sequencing depth leads the cell state and predicts the response to thinning

A description fitted to a model’s outputs can be tested by changing the input. We thinned the visible counts of every held-out test cell binomially to one half or one quarter, kept every masked query and its target, and asked the white box, fitted on untouched training cells, to predict how the Transformer’s outputs change (Fig. 4a). Thinning is a natural test of this description because it lowers the depth of a cell, its leading cell-level scalar, while the cell and the questions asked about it stay the same. Thinning moves the rank-target Transformer’s predictions by about three quarters of the spread of its whole contextual function at half depth. The white box predicts these changes across 2.0 million masked queries with *R*^2^ = 0.78 at half depth and 0.81 at quarter depth, and it does so in every one of ten cohorts (Fig. 4c,e).

**Figure 4.**
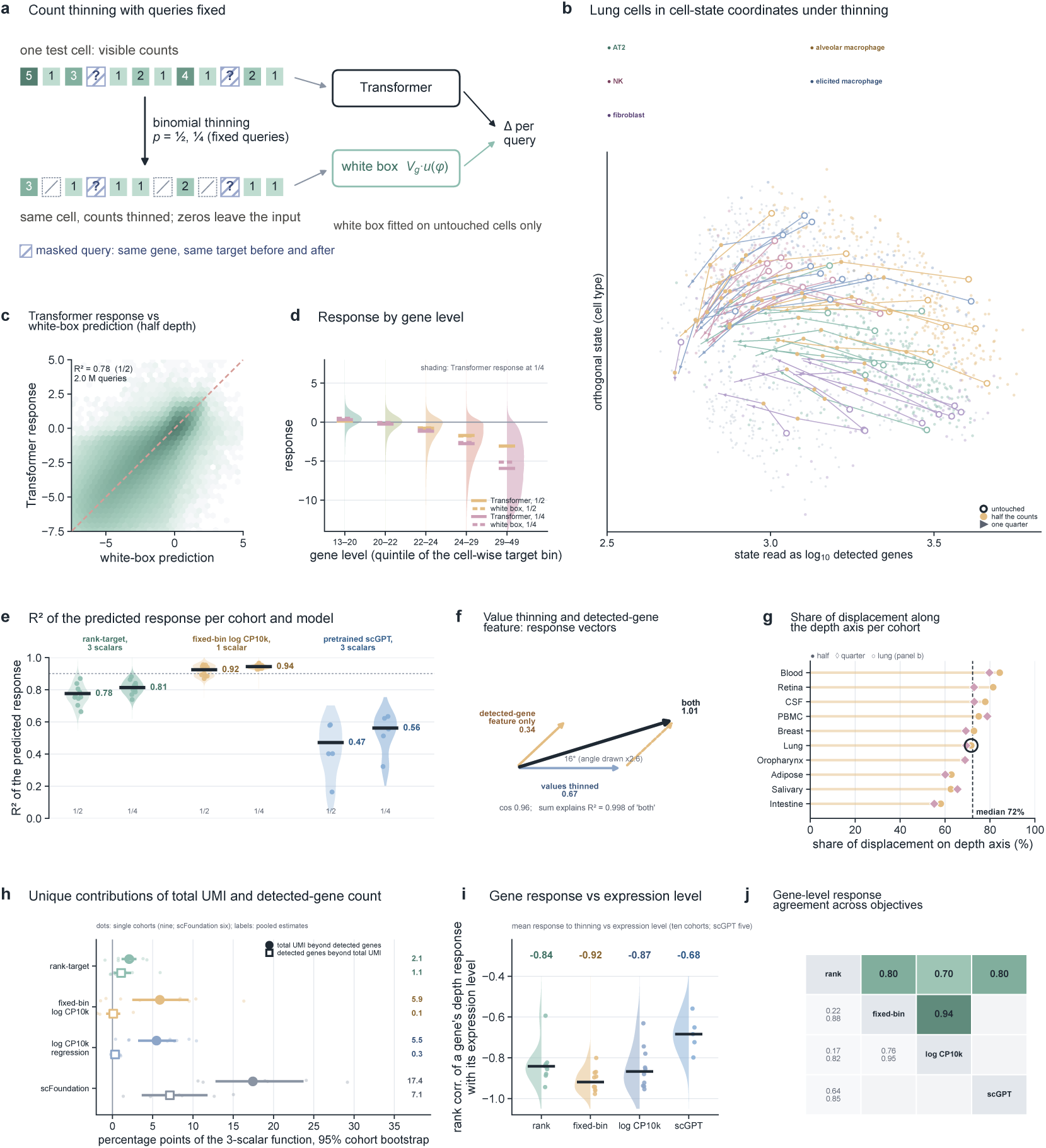
The white box predicts responses to count thinning along the depth axis. **a**, The schematic defines count thinning, binomial thinning of visible counts to one half or one quarter, with queries and targets fixed. Crossed boxes denote lost detections; arrows connect the rebuilt inputs to model and white-box responses (Methods). **b**, Lung-cell trajectories compare untouched and thinned cell states. Colours mark cell types; rings mark untouched cells, yellow dots half-depth states and arrowheads quarter-depth states for ten cells per type. **c**, Predicted and observed query responses are compared at half depth. Hexagon intensity shows query density on a logarithmic scale; the dashed diagonal marks equality. **d**, Responses are grouped by target-bin quintile. Shading shows quarter-depth response distributions; solid and dashed ticks mark model and white-box means at each dose. **e**, Response agreement compares rank-target, fixed-bin log counts per ten thousand (CP10k) and pretrained scGPT models. Dots denote cohorts, densities show their distributions, and black ticks mark pooled uncentred agreement. **f**, Separate value and detected-gene-feature interventions are compared with their joint response. Vectors show relative response lengths and their sum; vertical separation is exaggerated as labelled. **g**, Cell-state displacement is projected onto the depth direction across ten cohorts. Circles and diamonds mark half and quarter depth; the dashed line marks the half-depth cohort median and the ring identifies lung. **h**, Depth covariates compare objectives and inputs. Circles show total unique molecular identifiers (UMIs) beyond detected-gene count, open squares the converse, and bars 95% cohort-bootstrap intervals. **i**, Gene-response correlations with expression level compare objectives. Dots denote cohorts and black ticks medians. **j**, Gene responses are compared between objectives and scGPT. Upper-triangle shading encodes cohort-median correlations; lower entries give cohort minima and maxima.

Thinning moves cells along the depth axis of the cell state. That axis carries 72% of the squared displacement in the median cohort, and cells of every type move the same way. Genes answer according to their level: lowly expressed genes rise and highly expressed genes fall, and the white box follows both the ordering and the size of the mean response (Fig. 4b,d,g).

Which depth statistic leads depends on the objective and on how the inputs are built (Fig. 4h). The fixed-bin Transformer is trained on global bins of log CP10k, the logarithm of one plus counts per 10,000. In it, one cell-level scalar carries 99% of the white-box gain, against 75% under rank targets, this scalar follows total UMI, and a one-scalar white box predicts its response to thinning with *R*^2^ above 0.9. Total UMI leads because the fixed-bin inputs were normalized by full-cell totals; when the training inputs are rebuilt from visible genes alone, detected-gene count leads as closely. Under rank targets the two statistics are interchangeable, and in the one cohort whose counts were pre-normalized to a near-constant total, the function follows detected-gene count (Extended Data Fig. 8a,b). scFoundation orders the depth covariates in the same way, with cell type carrying as much. Pretrained scGPT’s response to thinning is predicted less completely (*R*^2^ = 0.47 and 0.56); 57–63% of the response lies within the span of its gene readouts (Fig. 4e). Fixed gene offsets cancel in this within-gene response and cannot explain the lower agreement.

Changing the values and changing a supplied depth feature move predictions along one direction. The log CP10k regression Transformer receives depth twice, through the visible values and through a detected-gene feature in its classification token. Thinning the values with that feature held fixed and changing the feature with the values held fixed give nearly parallel responses (uncentred cosine 0.96), and their sum predicts the joint change almost exactly (Fig. 4f). Across objectives and in scGPT, a gene’s response to thinning tracks how lowly it is expressed, and gene-level responses agree between objectives and with scGPT, even though the mean response changes sign between rank and log CP10k targets (Fig. 4i,j and Extended Data Fig. 8c).

Both targets make depth the best single summary of a cell. Library-size normalization places every single-count gene at log(1 + 10^4^*/*UMI), and ranking moves a gene’s target with the number of genes that compete with it. Either way, the best prediction of a masked gene depends on the cell’s depth, which the model reads from its visible genes and from any supplied depth feature, and coarse lineage supplies most of the rest of the cell state.

### Interpreting readouts requires matched functional controls

If masked prediction is a readout of a few cell-level scalars led by depth, the representations that interpretability methods read should carry the same statistics. We applied six common readouts to pretrained scGPT, to the controlled Transformers, to control models in which one input channel carries no information by construction, and to a 100,000-cell synthetic benchmark with 12 planted modules and regulators, each against a matched control. The cell state shows the distinction most directly. A linear probe decodes it from every layer in every cohort (*R*^2^ = 0.87 at layer 5), yet swapping it between cells along the probe directions transfers 0.23% of the output difference, less than swapping a random three-dimensional subspace (Fig. 5a–c). Learned subspaces can transfer more than the full state at late layers, but randomly initialized networks show the same effect (Extended Data Fig. 9b,i).

**Figure 5.**
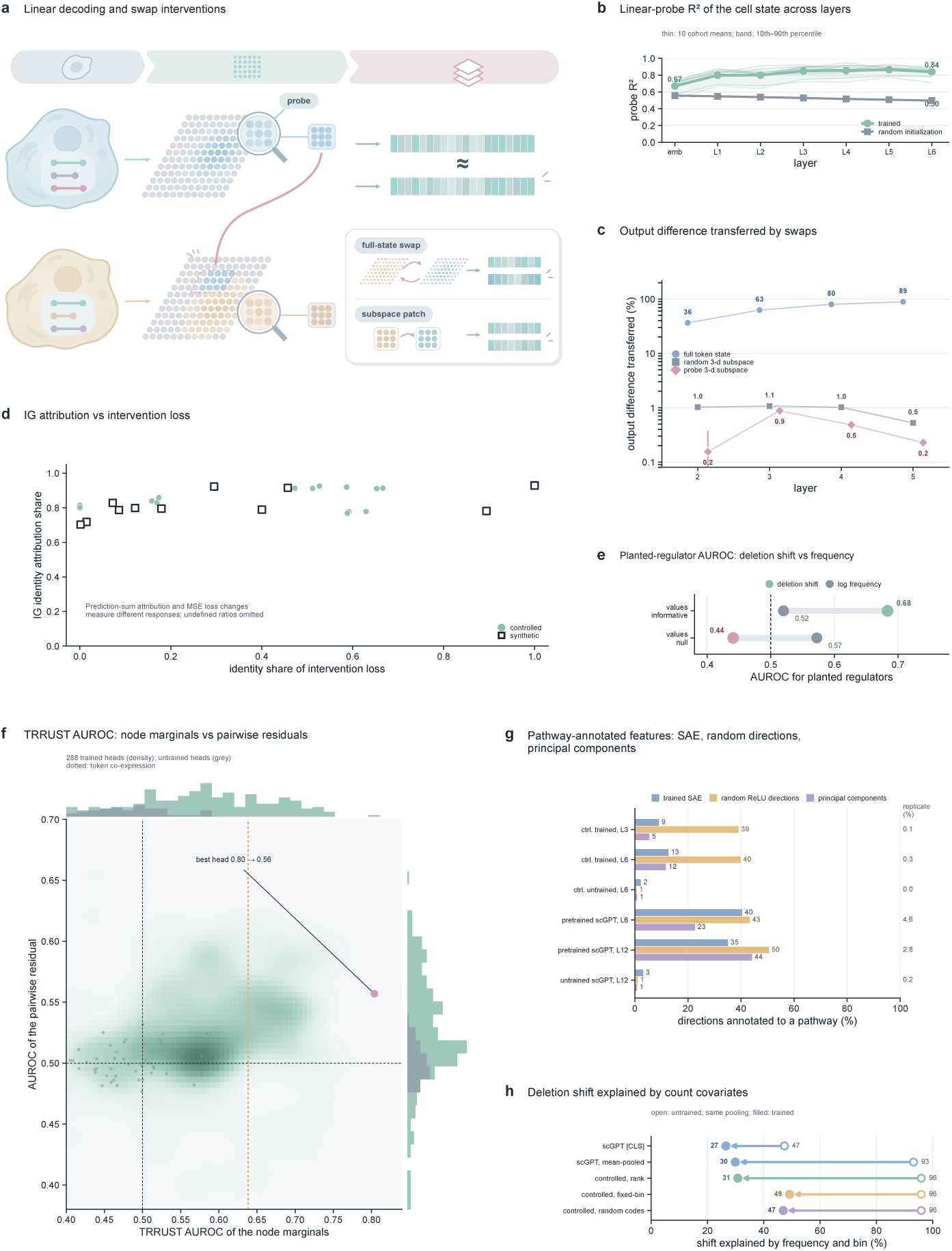
Matched controls distinguish readable structure from intervention effects. **a**, The schematic defines linear decoding and swap interventions on the same layer state. Branches distinguish reading a covariate from replacing its coordinates along probe axes. **b**, Linear probes decode cell state across layers of trained and randomly initialised Transformers. Thin lines denote ten cohort means across runs, thick lines the stored macro summaries, and shading the tenth-to-ninetieth cohort percentiles of those means. **c**, Swap interventions compare full-token, random-subspace and probe-subspace transfer. Symbols identify interventions; bars show the visible portions of 95% bootstrap intervals on a logarithmic transfer axis (Methods). **d**, Integrated Gradients (IG) identity attribution is compared with intervention-based identity loss share. Circles denote controlled-model runs and open squares synthetic runs, each averaged over available cohorts. The horizontal axis is the positive-part MSE increase from within-cell identity permutation divided by its sum with the increase from constant-value input. Zero or float32-resolution denominators are undefined and omitted. IG attributes the sum of predictions along embedding paths; no equality with this loss ratio is assumed. **e**, Deletion shifts and gene frequency are compared for recovering planted regulators. Points show area under the receiver operating characteristic curve (AUROC) for value-informative and value-null conditions; connectors pair readouts. **f**, Attention-head node marginals and pairwise residuals are scored against TRRUST, a curated database of human transcriptional regulatory interactions. Density represents 288 trained heads, grey dots untrained heads, marginal histograms their distributions, dashed lines chance, and the dotted line token co-expression. **g**, Pathway annotation compares sparse autoencoder (SAE) atoms with random rectified directions and principal components. Bars show annotated shares, averaging two SAE fits; right-hand labels show atom reproducibility at cosine similarity above 0.9. **h**, Count covariates predict deletion-induced embedding shifts in trained and untrained models. Open and filled circles distinguish training state, arrows pair matched pooling conditions, and position gives variance explained by frequency, expression bin and visible-gene count.

Cell embeddings carry depth. A polynomial in detected-gene count and total UMI explains about half of the embedding variance of the rank-target Transformer and most of it under fixed bins. A principal-component projection of the binary detection pattern, which uses no network at all, annotates cells better than these embeddings, and untrained Transformers match a random projection of the same pattern (Extended Data Fig. 9a,d). Pretrained scGPT is less dominated by depth and annotates better than the detection projection on the same cohorts. In the controlled models, these comparisons show that annotation can largely reflect summaries of gene detection and sequencing depth.

Integrated Gradients (IG) measures path-dependent attribution in the embedding space. For identity and value vectors added at the input, the path gradient is shared and IG multiplies it by each vector’s displacement from the baseline. Controls that randomise gene identities on every forward pass still receive 75–95% identity attribution. Across controlled and synthetic models, these attribution shares also differ from the relative loss increases caused by within-cell identity permutation and value clamping (Fig. 5d and Extended Data Fig. 9e). The two quantities measure different objects: attribution to a sum of masked predictions along an embedding path versus changes in prediction error after discrete input interventions. The IG channel share therefore cannot be read directly as the share of informative identity or value input.

Attention benchmarks reward hub genes. Decomposing each head’s attention into node marginals and a pairwise residual, the marginals of single heads separate 715 TRRUST regulator–target pairs from frequency-matched negatives with an AUROC of 0.80, while the pairwise residual stays near chance (Fig. 5f). Head-averaged attention is matched by Transformers with random gene codes and by token co-expression, and the two genes’ TRRUST degrees alone separate the pairs almost perfectly. Pathway benchmarks reward annotation degree in the same way (Extended Data Fig. 9c,h). Enhancer–promoter benchmarks behave alike: inverse distance reaches an AUROC of 0.88 on 14,734 CRISPRi-tested pairs, as high as the token similarity of a chromatin-accessibility model, and both fall to chance once negatives are matched for distance (Methods).

Deleting a gene moves a cell according to the gene’s frequency. Gene frequency, expression bin and the number of visible genes explain almost all of the embedding shift caused by a deletion in untrained networks, and a quarter to a half after training. Deletion still carries signal beyond frequency: on the synthetic benchmark it ranks planted regulators above frequency alone when expression values carry information, and below it when they do not (Fig. 5e,h).

Dictionary and gene-table readouts are matched by simple baselines. Random rectified directions enrich Gene Ontology and Reactome terms as often as trained TopK sparse autoencoders, or more often, and fewer than 5% of autoencoder atoms reproduce across fits (Fig. 5g and Extended Data Fig. 9f). The gene tables of the controlled Transformers stay close to their random initialization (Extended Data Fig. 10d), and counterfactual decomposition and steering do isolate latents whose effect depends on expression magnitude (Extended Data Fig. 10m,n). The same entanglement appears in chromatin accessibility: in controlled scATAC autoencoders depth can be decoded from the latent space, and an autoencoder trained to remove it loses most of its annotation as well (Extended Data Fig. 10p).

## Discussion

The models tested here can process expression values through a narrow pathway while retaining a wider backbone to combine genes. Whole-network narrowing has a different effect because it also constrains that backbone. These results motivate allocating width by computational role. The two-feature encoder provides a concrete design that improves the tested training and annotation outcomes, while the sparse-panel experiments identify a setting in which pairwise attention adds about 17 percentage points of prediction gain over set pooling.

Gene detection carries most downstream transfer gain in the evaluated annotation and surface-protein tasks. This connects the models’ input dependence to the discreteness of the measurements: many detected transcripts have the same small count, while detection itself varies across cell types and conditions. Magnitudes still contribute to some perturbation readouts. Evaluating encoding choices by task therefore gives a clearer account than assuming that any preserved transfer score requires precise expression values.

The white box identifies a shared functional component beyond cohort means: gene offsets and a gene-specific readout of a low-dimensional cell state. Depth leads in the controlled models, while lineage contributes more in scGPT. The dependence follows the objective and preprocessing, and the description predicts responses to thinning. Longer training adds dimensions, connecting training dynamics to the complexity of the prediction function. Future evaluations should measure how much a model adds beyond baselines built from depth and cell type, alongside its expression-prediction accuracy.

Interpretability requires a separate comparison between what can be decoded and what contributes through a proposed mechanism. Detection projections, gene-frequency controls, attention marginals, random directions and untrained networks provide accessible baselines for that comparison. Channel-uninformative models show that embedding-path attribution need not track information carried by an input channel. These controls complement observations that attention can recover curated regulatory edges [3] while adding little over gene-level features in perturbation-target prediction [22].

Depth also reflects biological variation: mitochondrial and ribosomal fractions couple measurement quality to metabolic and translational state. Distinguishing regulatory mechanisms from shared cell state calls for interventional data assessed against calibrated noise [10–12, 29, 30]. Future work should test how training to convergence changes the cell state and whether interventional objectives reveal regulation beyond depth and lineage. Output approximation and probe-subspace swaps constrain specific functional accounts; they do not establish a unique internal representation. The parameter savings also require direct throughput and memory measurements before being treated as deployment savings.

## Methods

### Cohorts, splitting and normalisation

RNA-based evaluation covered 18 human single-cell cohorts spanning three assay modalities (Extended Data Fig. 1a,d). Ten multi-donor droplet single-cell or single-nucleus RNA-sequencing studies came from CZ CELLxGENE Discover [31]. They cover peripheral blood after SARS-CoV-2 infection and mRNA vaccination [32], integrated lung core [33], subcutaneous white adipose tissue [34], oropharyngeal squamous cell carcinoma [35] and minor salivary gland in Sjögren disease [36]. They also include paediatric ileal lamina propria [37], primary breast tumour [38], autoimmune and immunodeficiency peripheral blood [39], developing fetal retina [40] and leptomeningeal cerebrospinal fluid immune cells [41].

Four multi-donor cellular indexing of transcriptomes and epitopes by sequencing (CITE-seq) studies [42] from NCBI Gene Expression Omnibus cover bone marrow mononuclear cells, trisomy-8 autoinflammatory peripheral blood, lecanemab amyloid-related imaging abnormality peripheral blood [43] and inflammatory bowel disease intestinal mucosa. Pooled CRISPR-interference (CRISPRi) Perturb-seq covers Jurkat and HepG2 [30], K562 and RPE1 [29]. Repository accessions and dataset identifiers are listed in Data availability; Extended Data Fig. 1 summarises cohort sizes, task assignments and preprocessing.

We evaluated public pretrained scGPT whole-human (33 million cells, 12 layers, width 512) [1], scFoundation 50M-0.1B-res (50 million cells, 19,264 genes) [2], Geneformer-V2 gc104M (12 layers, width 768, 4,096-token context) [3, 4], and Arc State/State embedding model SE-100M (State) (8 layers, width 1,024, 2,048-slot context) [5]. State initialised gene features with pretrained 5,120-dimensional ESM-2 protein embeddings [44].

Tissue and CITE-seq fitting, validation and evaluation used disjoint biological donors. Perturb-seq used perturbation split halves. Downstream normalisations, ranks, quantile boundaries and depth summaries used unmasked transcripts. Exceptions were the positive-only scFoundation protocol, which retained pre-masking depth, and scGPT calibration with full-cell quantile boundaries, both checked against visible-only recalculation below. Controlled pretraining computed per-cell summaries before query masking; evaluation rebuilt them from visible genes. The visible-input pretraining controls below test this difference.

For each CITE-seq antibody, we removed its encoding transcript and curated paralogs, receptor subunits and binding partners before computing visible counts per ten thousand (CP10k_vis_). Perturb-seq evaluation used 27,887 split-half cells from 966,728 source cells. We removed all in-vocabulary perturbation targets identically from perturbed and non-targeting cells before normalisation: 77 to 78 of the 80 targets per cohort, 310 total. Expression-regression inputs excluded the 500 target highly variable genes.

### Pretrained encoders, interventions and probes

Representations were extracted with dropout disabled (Figs. 1b and 2, and Extended Data Figs. 2 and 4). We centred each scalar value-embedding curve across input levels and applied singular-value decomposition, then projected onto its first *k* ∈ {1, 2, 4, 8} directions. Comparators were the original encoder, an ordinary-least-squares affine value encoder and a constant, evaluated on lung tissue and K562 cells.

For scGPT, expressed genes were encoded in 50 within-cell quantile bins, up to 1,199 genes plus the classification token at 2.0. The affine value encoder replaced the post-normalisation value-encoder output over *v* ∈ [1, 49], leaving the classification token unchanged. Clamped input set every detected-gene value to 25.5; value shuffling permuted bins among detected genes. Identity permutation sampled vocabulary genes of matched count while retaining the value multiset. Cell representations were *L*_2_-normalised classification-token states of width 512. scFoundation received counts over 19,264 genes, with both depth tokens set to log_10_(*N*_vis_), and returned pooled 3,072-dimensional representations. The affine fit covered positive values *v* ∈ [0.05, 6.0]. Clamped input set positive values to 1.0 while retaining depth tokens; the fixed-depth control set both tokens to 3.5. Value shuffling permuted positive values within cells; identity permutation shuffled nonzero gene identities.

Geneformer-V2 inputs were visible CP10k counts divided by published gene-median factors and sorted, with up to 4,094 gene tokens between boundary tokens. Representations were *L*_2_-normalised, mean-pooled penultimate-layer states of width 768. Controls omitted median scaling, randomised detected-gene order for clamped input, reversed rank order, permuted identities or used random initialisation.

State mapped positive unique molecular identifier (UMI) counts to 19,790 protein-coding genes, transformed them through log(1 + UMI) to relative weights 100*w_g_*, and sorted them into 2,047 slots, padding with unexpressed genes. Its 10-bin soft-count embedding produced *L*_2_-normalised classification-token states of width 1,024. The affine fit covered *c* [0.015, 0.60]. Clamped input set positive counts to 1 before weighting; a fixed-cardinality control held positive slot weights constant across cells. Shuffling permuted weights among expressed slots.

Net gain is the improvement over the cohort-mean predictor: the score increase for higher-is-better measures or the error reduction for masked prediction. For altered input *A*, retention is [*G*(*A*) − *G*(*I*)]*/*[*G*(*O*) − *G*(*I*)], where *G* is net gain, *I* is identity-permuted input and *O* is original input.

Probes were refitted for each input. Cell-type annotation used multinomial logistic regression; expression features were standardised using fit cells and divided by 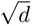, with regularisation selected on validation cells from {0.1, 0.5, 1, 5, 10, 50}. Label budgets were two labelled cells per class (two-shot), then 5, 10, 25, 50, 100 and 200 cells per class and the full split, with five subsampling replicates. Full-split annotation used a common five-cohort protocol; few-shot Geneformer-V2 and State scores used paired intervention runs. Temporal sensitivity excluded adipose tissue, leaving 19 of the 24 held-out donors.

CITE-seq prediction used donor-disjoint *L*_2_-regularised ridge regression of centred-log-ratio protein abundance, with donor-grouped cross-validation across 32 donors and 463 non-isotype channels, 461 with nonzero staining. Partial fine-tuning initialised linear heads from pretrained-representation probes [45], then trained the value module, top two Transformer blocks and readout using AdamW [46]. It covered two CITE-seq cohorts (4 held-out donors, 264 channels) and eight RNA-sequencing held-out donors evaluated at full labels and ten labels per class. Perturbation consistency was the cosine between uncentred shifts of *L*_2_-normalised embeddings, averaged over ten random split halves per perturbation.

The annotation fine-tuning comparison used adipose and oropharyngeal carcinoma donors, with full training splits and 10 labelled cells per class. Original, affine and clamped inputs were compared in scGPT and scFoundation, including clamped input with fixed depth. Its RNA-sequencing donors differ from the CITE-seq fine-tuning population; the latter supplies the protein comparison in Extended Data Fig. 4c.

### Chromatin-accessibility models

We analysed single-cell chromatin accessibility in bone marrow mononuclear cells (BMMC; GSE194122; 20,000 cells, 10 donors), brain (GSE312668; 18,000 cells, 9 donors), cerebellum (GSE331288; 21,660 cells, 10 donors) and DOGMA (GSE156478; 16,000 cells, 4 libraries). Three-fold grouped cross-validation used a stratified subsample of at most 2,100 cells per cohort, grouping by donor in BMMC, brain and cerebellum. DOGMA groups are its four libraries (LLL_CTRL, LLL_STIM, DIG_CTRL and DIG_STIM), not independent donors. Unlike the RNA transfer analyses above, each accessibility probe was fitted on intact training-fold embeddings and evaluated unchanged on test-fold interventions. Scores are macro-F1. We compared EpiAgent classification-token and mean embeddings, term frequency–inverse document frequency (TF-IDF) with latent semantic indexing, and ChromFound. Interventions binarised counts, permuted non-zero values within each cell or permuted peak coordinates within 20 cohort-level feature-detection-frequency strata using a fixed random seed. Paired BMMC and DOGMA RNA supplied scGPT controls.

The share of non-zero entries equal to one or two was computed on called peaks. Value-embedding effective rank was derived from singular values of the value or rank embedding (Extended Data Fig. 2k). Controlled BMMC models included a binary autoencoder, an autoencoder with a linear value input, a Poisson autoencoder with a depth adversary and a joint RNA–ATAC model, using 15,241 training cells, 4,759 held-out-donor test cells and three independent runs (Extended Data Fig. 10p).

### Count sparsity and detection-explained variance

We used raw UMI counts from 21,000 evaluation cells in adipose, oropharyngeal carcinoma, retina and cerebrospinal fluid [13, 14, 47, 48]. Salivary-gland pseudo-counts were excluded. Single- and double-molecule frequencies were computed over nonzero entries and averaged equally across cohorts. For gene *g*, *Y_g_* = log(1 + CP10k*_g_*) and *I_g_* = **1***{C_g_ >* 0*}*. Detection-explained variance was Var(E[*Y_g_ | I_g_*])*/* Var(*Y_g_*), from Var(*Y_g_*) = E[Var(*Y_g_ | I_g_*)] + Var(E[*Y_g_ | I_g_*]). We also computed it using held-out-donor cells only.

### Noise floors and masked-output calibration

Within-cell rank quantiles assign equal positive counts to a contiguous interval of integer bins {*a* + 1*, …, b*}. Uniform tie-breaking has conditional variance *σ*_tie_^2^ (*a, b*) = ((*b − a*)^2^ *−* 1)*/*12. Averaging over masked queries gives the tie floor *L*_tie_; subtracting it from the out-of-cohort gene-mean error *G* gives the post-tie window *W*_tie_. These are squared-bin quantities. We report cohort-equal, query-weighted and mean-of-cohort-ratio aggregations separately (Extended Data Fig. 5e).

For the technical-sampling sensitivity [47, 48], we held other counts fixed and resampled one positive masked query from a zero-truncated Poisson distribution, P(*C* = *k C >* 0; *λ*) = (*λ^k^/k*!)*/*(*e^λ^* 1). The upper truncation was 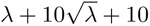, with tail mass below 10^−11^. Integrating tie-breaking over recomputed boundaries gave *L_cg_*^*^ = E[*m*_2_(*C_cg_*) *| C_cg_ >* 0] *−* (E[*µ*_bin_(*C_cg_*) *| C_cg_ >* 0])^2^, and *W^*^* = *G − L^*^*. Expression rates used empirical-Bayes neighbour smoothing with *k*_nn_ 10, 100 and 20 global-prior pseudo-counts. We also measured whether single-query resampling changed a visible bin, against a 5% threshold; this distinguishes the sampling assumption from the exact tie calculation.

Post-hoc scGPT prediction used 25% positive masked queries and 50-bin targets. scFoundation used either mixed queries (30% positive and 3% zero) with visible normalisation or positive-only queries (25% positive) retaining full-cell depth. Calibration was evaluated without target-cohort fitting and with 3,000 target-cohort training cells plus 750 validation cells, disjoint from held-out donors. For scGPT, candidate features combined expression-decoder and cell-context-decoder predictions, empirical-Bayes training gene means, gene- by-detection-count lookups, visible gene count and log visible transcript depth. Regularised per-gene and cross-validated ridge combinations fitted the target bins.

For scFoundation, calibration combined decoder predictions and detection-rate de-attenuation, pred*/P*_det_, with out-of-cohort gene means, visible detection and log visible depth. Detection rates came from out-of-cohort marginals or *k* = 100 binary-detection neighbours; empirical-Bayes per-gene ridge blending combined features. Marginal-only rescaling tested dependence on neighbour lookup. Positive-only scores and mixed-query scores use different targets and normalisations and are presented separately. scGPT boundary sensitivity recomputed quantiles using visible genes only before repeating calibration.

### Controlled pretraining and dimensionality

Controlled Transformers (Transformers trained from scratch under controlled conditions) used six pre-normalisation layers, width 256, 8 heads, up to 1,024 tokens, 20,080 protein-coding genes and AdamW [46], trained on 199,936 cells from training donors of all ten tissue cohorts. Three independent runs shared data order, masks and schedules across conditions. The rank-target objective used 50 within-cell rank bins. Visible tokens attended to visible keys; masked queries attended to visible keys and themselves, preventing other tokens from reading masked values.

The MLP value encoder was a two-layer rectified-linear network with layer normalisation. Its affine comparator was *a* + (*v/*50)*b*, *a, b* R^256^, variance-matched at initialisation. Learned gene tables were compared with random 32-dimensional Gaussian codes, row-normalised to length 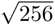 and projected through a shared orthogonally initialised 32 *×* 256 layer with layer normalisation. Both had trainable per-gene output biases.

The width scan used *d* ∈ {64, 128, 256, 512}, with the same construction scaled to *d* (Fig. 1g and Extended Data Fig. 2e,h). Random-code five-epoch runs used three replicates at every width; ten-epoch runs used three at width 256 and one elsewhere. Learned-code runs used three at width 256 and one at widths 64 and 512. At width 512, learning-rate multipliers 0.5*×*, 1*×* and 2*×* tested optimisation dependence.

Calibration used 3,000 training-donor cells from the target cohort and in-cohort gene means. Without target-cohort calibration, predictions were evaluated against out-of-cohort means.

Whole-network narrowing trained every component at *d* ∈ {512, 128, 64, 32, 16, 8, 4} for 10 epochs on 200,000 cells, with three runs. The two-track Transformer retained width 512 for identity, attention routing and cell embeddings, while values stayed at *d*_val_ ∈ {512, 64, 4} through every layer and the masked head. The value track contains 20,488,196 parameters at width 512 and 2,080 at width 4; total parameters are 35,500,554 and 15,014,438, respectively. Figure 1i retains component-specific denominators: whole-network backbone/value parameters for the whole-network scan and value-stream parameters for the two-track scan, each relative to its own width-512 reference. Value-track learning rates were 6 10^−4^ at width 512 and 3 10^−3^ at width 4, with 8 and 4 heads, respectively.

Longer narrow whole-network runs are shown in Extended Data Fig. 3c,d.

Width scans express masked-prediction net gain relative to the width-512 model of the same architecture. Donor-paired relative error was assessed over 59 donors with a 1% equivalence margin. Intervals in Extended Data Fig. 3a,b are two-sided 90% Student’s t intervals on paired donor differences. Annotation used linear probes on 24 donors from five cohorts and a separate 40-donor protocol, with a *±*1.5-point margin; the value-track comparison meets this margin on the latter protocol. Its 24-donor 90% interval is [*−*1.5077, +0.6051] points and narrowly fails equivalence; the 40-donor interval is [*−*0.7754, +1.0033] and passes.

The 15-arm value-encoder bottleneck sweep used three independent runs per arm, 200,000 cells, 20,080 genes and five epochs, at hidden width 512 (768 for the scFoundation-sized block). Blocks used rank-target or continuous log CP10k targets; bottleneck widths were *k* 1, 2, 4, 8, 16, 32 and full width, alongside an affine value encoder. Gates required masked-prediction mean squared error within 2.5%, net gain within 2.5 percentage points and annotation within 1 point of the block’s full-width encoder. Other downstream probes used 0.010 non-inferiority for expression and surface-protein correlation, overall and within cell type or lineage. Two-one-sided tests used *α* = 0.05 and 1,000 bootstrap draws. Effective rank counted singular directions of value-network output needed for 99% variance. The nonlinear K1 arm passes masked-prediction and downstream criteria but exceeds the upper gain-equivalence margin; failure of this gate is not evidence of insufficient capacity.

### Two-feature value encoder

The two-feature encoder used the rank-two affine map *a* + *q*_mid_*b*_rank_ + *z*_amp_*b*_amp_.

Here *q*_mid_ is the tie-free median rank of a count among the cell’s detected genes, and *z*_amp_ is log count standardised within cell. Raw depth was excluded from the classification token ([CLS]); the readout head was anchored to visible moments (mean, standard deviation, minimum positive value and log depth). The controlled six-layer, width-256 Transformer used 20,080 genes, three independent runs, five epochs, ten cohorts and 59 held-out donors. The two-feature encoder had 1,280 value-encoder parameters versus 66,816 for the nonlinear network. Mid-rank-only and amplitude-only arms each had 768 parameters; a depth-leak control supplied depth through [CLS]. A common continuous target supported donor-paired contrasts against quantile affine, mid-rank-only and other ablation arms.

The graft replaced pretrained scGPT’s 264,704-parameter value encoder with 2,560 parameters, unfroze layers 8–11 and trained for two epochs (3,124 steps) on a donor-disjoint 200,000-cell corpus. Scaling used a 12-layer, width-512 model and 1,000,000 cells.

### Sparse-panel predictors and attention controls

The controlled 1,024-gene positive-expression panel used donor-disjoint peripheral blood, 1,999 test cells and 5,230 queries, averaging 17.96 visible genes per cell. Gain share compared the training gene mean with the mean risk of the three reference Transformers trained for 12 passes. Non-neural predictors received visible identities and bins plus masked query identities. They combined detected-count-conditioned Gaussian means, gradient-boosted closure trees over visible and query summaries, and binary-detection Jaccard neighbours (*k* = 200); incremental contributions were averaged over both orderings.

Architecture-matched Deep Sets [49] pooled visible genes and masked identities without self-attention. They and Transformers followed a 48-pass cosine schedule, evaluated at passes 12, 24 and 48. Further controls pooled visible genes alone, blocked masked keys, or kept masked keys and values at normalised initial embeddings. Positive-only panel queries reveal detection through their identity; full-vocabulary retraining tested the consequence.

Depth variants used L ∈ {1, 3, 5} visible-context layers before one readout layer or one self-attention layer. Mask-routing controls removed mask-to-visible or mask-to-mask attention, either post hoc or during retraining (Extended Data Fig. 7).

Post-hoc routing interventions removed masked and off-diagonal visible edges from already trained panel models. Retrained controls used the identical attention mask during fitting. The mask-ratio comparison included updated attention, static masked keys and values, retrained sparsity and set pooling. Error differences between set pooling and attention were tabulated jointly by masked-query count and visible-gene count; cells with fewer than 100 queries were omitted. Further summaries grouped queries by masked-gene count or visible-count quartile, and repeated the comparison for query UMI at least 6 (Extended Data Fig. 7e–g,j,k).

Panel white-box fits compared exact alternating least squares with an Adam joint fit across numbers of cell-level scalars. Gain share and distillation agreement used the same panel reference and Transformer outputs. Multiple random initialisations and initialisation from the Adam solution checked sensitivity to starting state. Full-context subsampling removes visible evidence from an existing full-vocabulary task; the sparse positive panel changes the vocabulary and restricts queries to detected genes.

### Prediction score and function agreement

For Transformer *T*, predictor *S* and reference *P*, distillation *R*^2^ = 1 ∑(*T − S*)^2^*/* ∑(*T − P*)^2^ over test queries. Gain share is [E(*Y − P*)^2^ E(*Y S*)^2^]*/*[E(*Y − P*)^2^ E(*Y − T*)^2^]. If *T* and *S* are conditional expectations under nested information, with *S* = E[*T* information of *S*] and *P* nested in *S*, squared-error orthogonality makes the two quantities equal. Calibration was checked through gain divided by deviation energy, which equals one for a calibrated predictor. Full-vocabulary retrieval is used to compare score and output agreement, without assuming matched information.

### White box and cell covariates

The reference Transformer *T* averaged three controlled full-vocabulary Transformers with learned gene codes and affine values, evaluated after five epochs of a ten-epoch schedule. Fitting used training donors within each cohort; evaluation used held-out donors. We define the function as the model’s function beyond cohort means. The reference *P_g_* was the fit-cell mean bin, shrunk with five pseudo-counts toward the cohort-wide mean.

The low-rank linear white box was 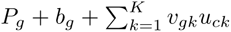. Alternating least squares fitted free cell states and gene readouts to Transformer outputs, minimising squared error plus *λ*(∥*U*∥ ^2^ + ∥*V*∥^2^), with *λ* = 3, a truncated singular-value initialisation and at most 40 iterations. Linear-kernel ridge regression then mapped visible features to cell state, *u_c_* = *Wϕ*(*c*). Five-fold cross-validation on fit cells selected ridge penalty from *{*0.1, 1, 10, 10^2^, 10^3^, 10^4^*}* by query-level prediction error.

Features comprised visible binary detection over the vocabulary, standardised visible bins (*q −* 25.5)*/*15, and four summaries: log(1+*n*_vis_), bin mean, bin standard deviation and fraction of bins at least 40. Test predictions used only these features. Cohort results pooled sums of squared errors. Offsets alone corresponded to *K* = 0; shuffling permuted test features within cohorts. Reproducibility compared each run with the mean of the other two. Pretrained scGPT used the same fitting procedure on its own visible genes and bins in five donor-disjoint cohorts.

Covariate fits replaced *ϕ* while retaining gene readouts. Covariates were whole-cell log detected-gene count log *D*, one-hot coarse cell type, visible summaries, log total UMI, mitochondrial fraction and ribosomal fraction. Single covariates used *K* = 1 and combinations *K* = 3. The share of the white-box increment was 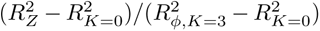; the fraction of the full function beyond offsets was 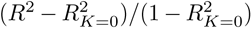.

For enrichment, cohort gene readouts were stacked, reduced to three pooled singular vectors and varimax-rotated. Additional axes projected gene–depth correlation, cell-type effects orthogonal to depth and the remaining direction onto the same span. Scores were tested raw and after regression on mean bin, detection rate, bin standard deviation and log mean count. The 1,801 gene sets included Hallmark, Reactome sets with at least five vocabulary genes and curated quality, stress, cell-cycle and lineage programmes. Weighted running-sum tests used 150 random sets per logarithmic size bin and Benjamini–Hochberg correction within axis.

The free-state control retained fit-cell gene readouts and fitted each test state on half its queries, scoring the other half and reversing the split. Single-model reliability was the uncentred correlation of independent deviations from *P*. Sparse-context reliability used the Spearman–Brown correction *r*_3_ = 3*r*_1_*/*(1 + 2*r*_1_).

### Rank arithmetic and state geometry

Gene depth responses were slopes of Transformer deviations or targets on standardised log *D*. A multinomial simulation used each cell’s total UMI and fit-cohort mean gene proportions, with three draws per cell, 50-bin quantiles, uniform tie-breaking, a 1,024-gene window and 15% masking. We correlated simulated target slopes with Transformer slopes across cohort–gene pairs. Ridge regression of log *D* on visible features measured window information about whole-cell detection.

For state plots, if *w_g_* is gene *g*’s query share and 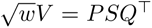, coordinates *SQ*^T^*u_c_* have length equal to the prediction’s root-mean-square change. Rotation aligned the first axis with least-squares log *D* and the second with the largest orthogonal direction; cell-type separation used *η*^2^. Query-level count, bin, prediction and depth summaries were averaged over depth deciles. Window-ranked controls recomputed targets among the same selected window genes, retaining architecture, masks, order and schedule.

### Thinning and objective controls

Count thinning (binomial thinning of visible counts to one half or one quarter) used 1,500 held-out cells per cohort. Zeroed genes left the input and visible bins were recomputed. Queries and targets stayed fixed. A white box fitted to untouched training cells predicted each response as 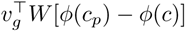.

Agreement used uncentred response *R*^2^ pooled across cohorts. The response fraction within gene-readout space used split-half free states. The intervention jointly changes detected-gene count, library size and bins. The depth direction was the least-squares map from untouched test states to log_10_ *D* within cohort. Its displacement share was summed squared projection divided by total squared displacement; we report the cohort median. The thinned whole-cell detection estimate scaled the original count by the surviving fraction of visible genes. Pretrained scGPT used visible-only quantiles, unchanged queries and matched tie-breaking draws, with its own fit-cell white box and target means.

Objective controls trained the same six-layer width-256 Transformer on 200,000 cells for five epochs with three replicates. Fixed-bin log CP10k targets used full-cell log CP10k assigned to 50 global bins, with edges at the 2nd to 98th percentiles of a 20,000-cell subsample. Visible values used those bins. After thinning, values were rebuilt using visible library size scaled by the retained count share and the same global edges. Auxiliary target readouts used stop-gradient hidden states.

The continuous log CP10k model used affine values rescaled within the visible set and two logarithmic features projected into the classification token: detected-gene count and visible CP10k sum. During pretraining the detected-gene feature used 0.85 times whole-cell detection; evaluation used visible detection. In separate interventions we thinned values with this feature fixed, changed only this feature, or changed both. The CP10k-sum feature followed the values. Pretraining bins and normalisation otherwise used full-cell summaries before masking. Visible-input controls recomputed rank and fixed-bin inputs after masking during training, retaining targets, masks, order and schedule.

Total UMI was recovered as the median of *c* 10^4^*/*(*e^y^*-1) across detected genes, where *c* was raw count and *y* stored log CP10k. Salivary-gland pseudo-count totals carry little library-size information and were excluded from comparisons of UMI and detection, with separate reporting. Unique contributions were the increment from the two-covariate fit over each single-covariate fit, relative to the white-box increment.

scFoundation held out 30% of detected genes and 3% of measured zeros, using visible-library log CP10k and two visible-depth tokens. Fits and tests each used 1,500 cells from seven cohorts; depth comparisons excluded salivary gland. Positive-query targets retained full-cell normalisation. Visible features comprised detection, standardised values and log visible library size. White-box, covariate and free-state analyses followed the procedures above; a sensitivity analysis included sampled zeros.

### Sparse context, scale and training duration

Context subsampling retained *n* ∈ {64, 128, 256} random visible genes and refitted the white box using retained features. An in-distribution control trained three models with a 64-gene visible cap, preserving queries and replacing dropped tokens with padding identities.

Scaling used a 12-layer width-512 Transformer with 8 heads and the same target, window and mask. Its 1,000,000-cell corpus included the 200,000-cell corpus plus 29 external datasets, excluding evaluation donors. Checkpoints after 7,812 steps (one pass, three runs) and 23,436 steps (three passes) were compared with the smaller corpus at matched steps. One-pass controls annealed at 7,812 steps; a second three-pass run and ten-epoch controlled checkpoints tested training duration. The earlier affine continual-adaptation comparison trained scGPT’s value encoder, top four layers and readout for two epochs, retaining lower layers and the gene table.

The one-pass million-cell checkpoint matches five passes over the controlled corpus in optimisation steps. Annealed one-pass controls separate stopping time from final learning rate. Three-pass model agreement compares nonlinear and affine encoders; controlled-model agreement compares independent runs of the same configuration (Fig. 3h and Extended Data Fig. 7a).

It complements the visible-input encoder and independent-model agreement.

### Interpretability readouts and controls

Integrated Gradients [50] in the primary additive-embedding and synthetic audits used 24 midpoint integration steps; the additional channel-control runs used 16. Baselines were zero vectors, a mean identity embedding over the evaluation batch with a mid-range value embedding, and independently shuffled visible identities and values. The scalar endpoint was the sum of masked predictions. Identity share was visible-identity *L*_1_ attribution divided by visible identity-plus-value *L*_1_ attribution; query-identity attribution was stored separately. Added identity and value vectors receive the same path gradient, multiplied by their respective displacements from baseline. Channel-uninformative controls randomised identities on every forward pass or clamped values before encoding. IG varied embeddings downstream of those operations and therefore evaluated paths that the discrete preprocessing need not produce; separate step-sensitivity checks varied integration step counts across baselines.

Within-cell identity permutation and value shuffling are inverse permutations, so their expected losses coincide for order-invariant readouts. Their loss ratio cannot identify channel use. Figure 5d reports the identity-permutation versus clamped-value loss ratio, 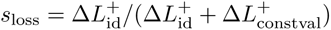, where *x*^+^ = max(*x,* 0). This is an intervention-loss summary, not a causal truth for IG. A zero denominator, or one no larger than a single float32 unit in the last place at the largest of the three losses, is treated as numerically undefined and omitted from ratio plots. The latter rule removes one clamped-value-control ratio built from a loss change of 0.00001526.

Embedding analyses used ten donor-disjoint cohorts for controlled models and five for scGPT. Depth was a polynomial in log detection and log UMI; annotation used standardised linear probes. For scGPT, five-fold cross-fitting estimated covariate-explained variance at every layer, and donor-grouped ridge classification assessed annotation before and after training-fold depth removal.

Deletion analyses used 19,200 controlled-model deletions and 4,000 scGPT deletions, trained and untrained, with classification-token and mean pooling. Nuisance regression predicted embedding-shift norm from log gene frequency, bin and visible-gene count. Attention over 2,400 cells and the 3,000 most expressed genes was decomposed into node marginals and residuals, *Āij* = *m_i_* + *m_j_* + *R_ij_*. TRRUST, a curated database of human transcriptional regulatory interactions, supplied 715 positive pairs and frequency-matched negatives. We reported head-averaged and per-head area under the receiver operating characteristic curve (AUROC).

The real-data TopK sparse autoencoders used 1,024 features, 32 active features and two fits per layer, compared with 1,024 random unit directions passed through a rectified linear unit. Controlled fits used 2,500 Adam steps and scGPT fits 2,000, batch size 4,096 and learning rate 0.001. Inputs were centred per channel and divided by a single global standard deviation; decoder atoms were renormalised after every step. The synthetic-module audit instead used 512 features, 16 active features and one 2,000-step fit per model/layer, with matched 512 random directions. Gene-table controls included annotation-degree sum, gene frequency, five gene-level scalars and positive pointwise-mutual-information factorisation of binary co-occurrence. Extended Data Figs. 9 and 10 show probe, swap, dictionary and attribution controls across all evaluated conditions and readouts.

Swap interventions (interchange interventions) compared swapping the full visible-token state with swaps restricted to a random subspace or the linear probe directions for the white-box cell state, across layers 2–5. Output-difference transfer was compared with linear decoding accuracy, using the same target cell state. Learned subspaces were assessed in trained and randomly initialised networks; distributed alignment search was evaluated at dimensions *k* = 1, 3, 6 and layers 1, 2, 3 and 5. Random-subspace, mismatched-state and gene-split evaluations tested transfer specificity, with the last scored on held-out genes (Fig. 5c and Extended Data Figs. 9b,i and 10j,k). These controls distinguish the ability to read a state from the effect of replacing it along particular directions.

Dictionary comparisons evaluated controlled layers 3 and 6, pretrained scGPT layers 6 and 12, and untrained reference layers. Enrichment used the top genes of each feature against GO and Reactome terms, including frequency- and annotation-degree-matched shuffles. Decoder-atom reproducibility used cosine > 0.9 across independent fits. On the synthetic benchmark, feature recovery was the median across planted modules of the best single-feature AUROC; trained and untrained models were compared on value-informative and value-null data, with an AUROC 0.8 recovery threshold (Extended Data Fig. 9g).

Counterfactual dictionary readouts grouped active latents into support-dominant, amplitude-dominant and synergistic tiers. The evaluated models were pretrained scGPT, two-feature encoder scGPT with low-rank adaptation and the million-cell two-feature encoder model. Dose-response plots compared latent steering with mean s.d. for 16 random latents at *α* = 2 (Extended Data Fig. 10m,n). Gradient-times-input attribution stability compared two tie-breaking draws using Spearman correlation within UMI strata 1, 2 and at least 5, alongside top-50 Jaccard overlap (Extended Data Fig. 10f). These are separate evaluations from the synthetic-module recovery test.

The synthetic benchmark generated negative-binomial counts for 600 genes across six cell types, with 40 transcription factors and 12 modules of 40 non-transcription-factor genes. In the value-null condition, detection depended on activity, whereas positive counts depended only on gene baseline and depth. Models used 100,000 training cells, 6,000 test cells, two independent runs and five epochs. These conditions supplied planted regulators and modules for deletion and dictionary controls.

The enhancer–promoter benchmark used the EngreitzLab CRISPR_comparison EPCrisprBenchmark GRCh38 tables: 10,356 K562 pairs and 4,378 pairs from five held-out cell types, totalling 14,734 valid pairs with 661 positives. Labels were Regulated; rows required valid connections, finite coordinates and autosomal or sex-chromosome membership. Distance was the supplied distanceToTSS. We scored inverse distance and ChromFound token cosine by AUROC overall and after distance matching. Within each source panel, positives in a fixed random order were matched to up to five unused negatives within 0.05 log10-distance units (distance floored at 100 bp), falling back to 0.12 when no candidate existed. Positives without candidates were omitted; negatives were not reused.

### Statistical analysis

Donor-level annotation and CITE-seq contrasts used 10,000 stratified donor-bootstrap draws and paired two-sided sign-flip tests. Geneformer-V2 and State summary retentions and four-cohort Perturb-seq means were point estimates; scGPT and scFoundation donor-level comparisons had paired intervals. Multi-run retraining reports arithmetic mean and standard deviation of individual risks. Panel contrasts used 4,000 two-level run-by-cell bootstrap draws.

Unless specified otherwise, function-analysis intervals were 95% percentile intervals from 4,000 cohort resamples, recomputing pooled values with deviation-energy weights and pairing cohorts for contrasts; tie-breaking variance was evaluated directly by resampling without parametric count assumptions, alongside Poisson count-noise estimates.

## Data availability

The ten single-cell and single-nucleus RNA-sequencing tissue cohorts were obtained from CZ CELLxGENE Discover [31] under dataset identifiers 242c6e7f-9016-4048-af70-d631f5eea188 [32], 066943a2-fdac-4b29-b348-40cede398e4e [33], 9d8e5dca-03a3-457d-b7fb-844c75735c83 [34], b6b5ea88-e092-46e4-b9c5-e93b52d5c195 [35], df1edb87-e512-43ae-b5f4-cb179cfc2bb4 [36], 6ceeaa86-9ceb-4582-b390-6d4dd6ff0572 [37], 9fddb063-056d-4202-8b8a-4b0ee531d3ce [38], c2a461b1-0c15-4047-9fcb-1f966fe55100 [39], aecc2743-4ff2-4b55-ac8c-a1ff76fc4bb4 [40] and 0bf30b02-db97-4560-b9cd-0fe3588892af [41]. CITE-seq data are available from NCBI Gene Expression Omnibus (GEO) under accessions GSE194122, GSE272583, GSE316096 and GSE331003. The four Perturb-seq cohorts were obtained from the published Replogle et al. [29] and Nadig et al. [30] resources. Chromatin-accessibility and paired multiome data are available from GEO under accessions GSE194122, GSE312668, GSE331288 and GSE156478. The enhancer–promoter benchmark used the GRCh38 training-K562 and held-out-five-cell-type tables from the EngreitzLab CRISPR_comparison EPCrisprBenchmark resource, as specified in Methods.

Processed summary tables underlying Figs. 1–5 and Extended Data Figs. 1–10 are available in the public repository specified below. Original raw datasets remain available from their respective repositories.

## Code availability

Code for model training, evaluation, statistical analysis, synthetic-benchmark generation and figure reproduction is available under the MIT License at https://github.com/zhang-lecheng/scfm-value-dimensions.

## Acknowledgements

This work was supported by the National Science and Technology Innovation – Major Program (No. 2022ZD0115101), the National Science and Technology Major Project of China (No. 2021YFA1301603), the SOE Dean Special Project Fund (SOE-DSPF) Program of Westlake University (Project No. WU2025B006), and the Center of Synthetic Biology and Integrated Bioengineering of Westlake University (Project No. WU2024A004).

## Author contributions

L.Z. and Y.Y. contributed equally.

L.Z. conceived the study, designed and executed the experiments, analysed the data and wrote the initial draft of the manuscript.

Y.Y. contributed to experimental evaluation, technical validation and manuscript revision.

S.Z.L. supervised the project, provided guidance and revised the manuscript. All authors approved the final manuscript.

## Extended Data Figures

## Extended Data Figures

**Extended Data Figure 1.**
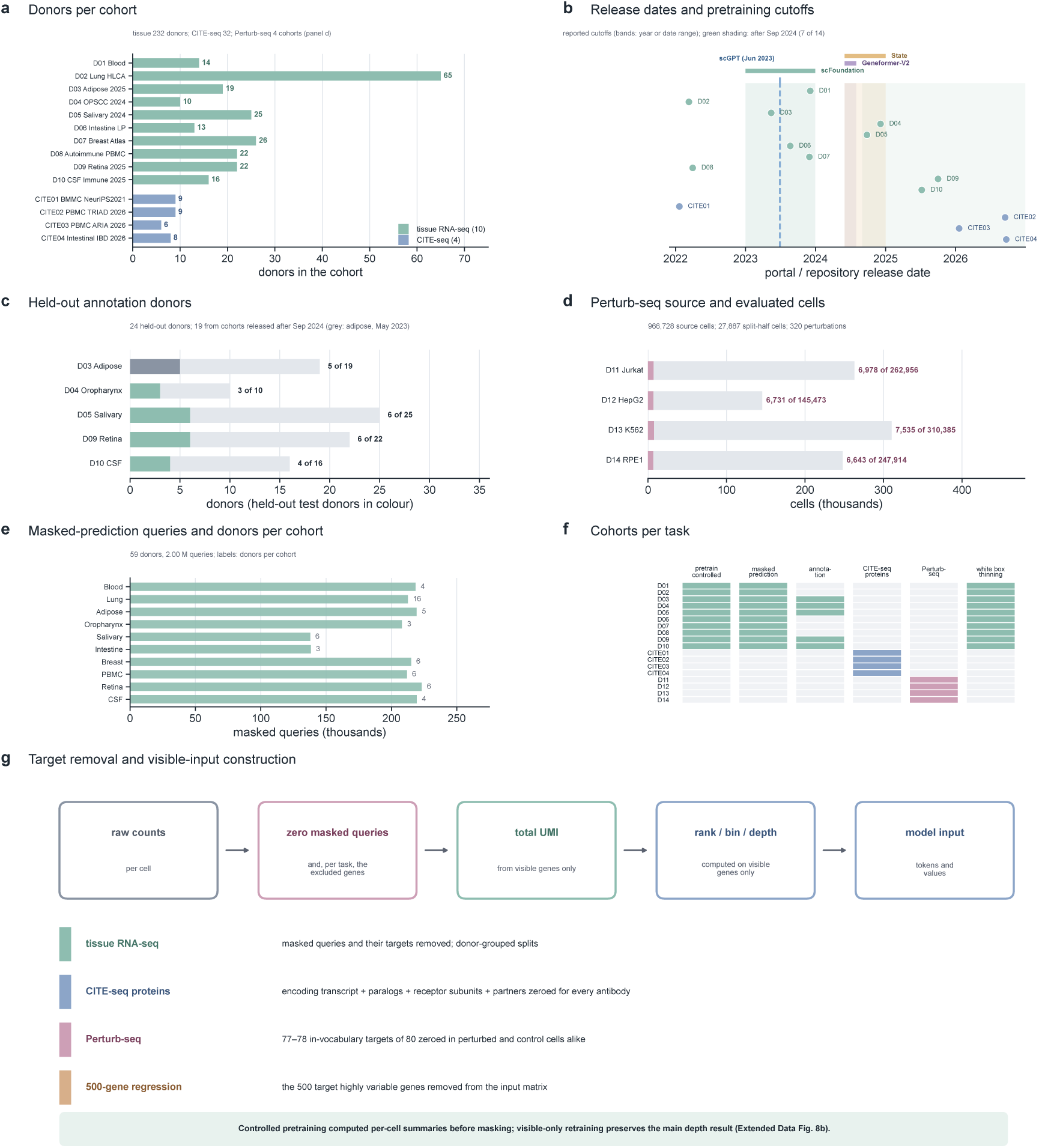
Cohort timing and donor splits define the evaluation populations. **a**, Cohort inventories compare tissue transcriptomes and paired transcript–surface-protein measurements. Bars show donor counts, colours distinguish modalities, and perturbation cohorts appear in d. **b**, Repository release dates are compared with model pretraining cutoffs. Points mark cohorts, lines mark dated cutoffs, bands mark cutoff windows, and green shading marks the later-release subset. **c**, Cell-type annotation cohorts show donor allocation. Coloured bars identify held-out donors within cohort totals; grey distinguishes adipose tissue in the temporal sensitivity analysis. **d**, Perturb-seq cohorts compare source and evaluated populations across 320 perturbations. Pale and coloured bars distinguish source and split-half evaluation cells, respectively. **e**, Masked-prediction cohorts show evaluation coverage. Bar lengths encode query counts and labels donor counts across 59 held-out donors. **f**, A matrix assigns cohorts to tasks. Coloured tiles mark participation; colours identify modalities. **g**, The schematic defines target removal and visible-input construction. Arrows order preprocessing, task-specific rows identify excluded transcripts, and the footer distinguishes controlled pretraining summaries from visible-only rebuilding (Methods).

**Extended Data Figure 2.**
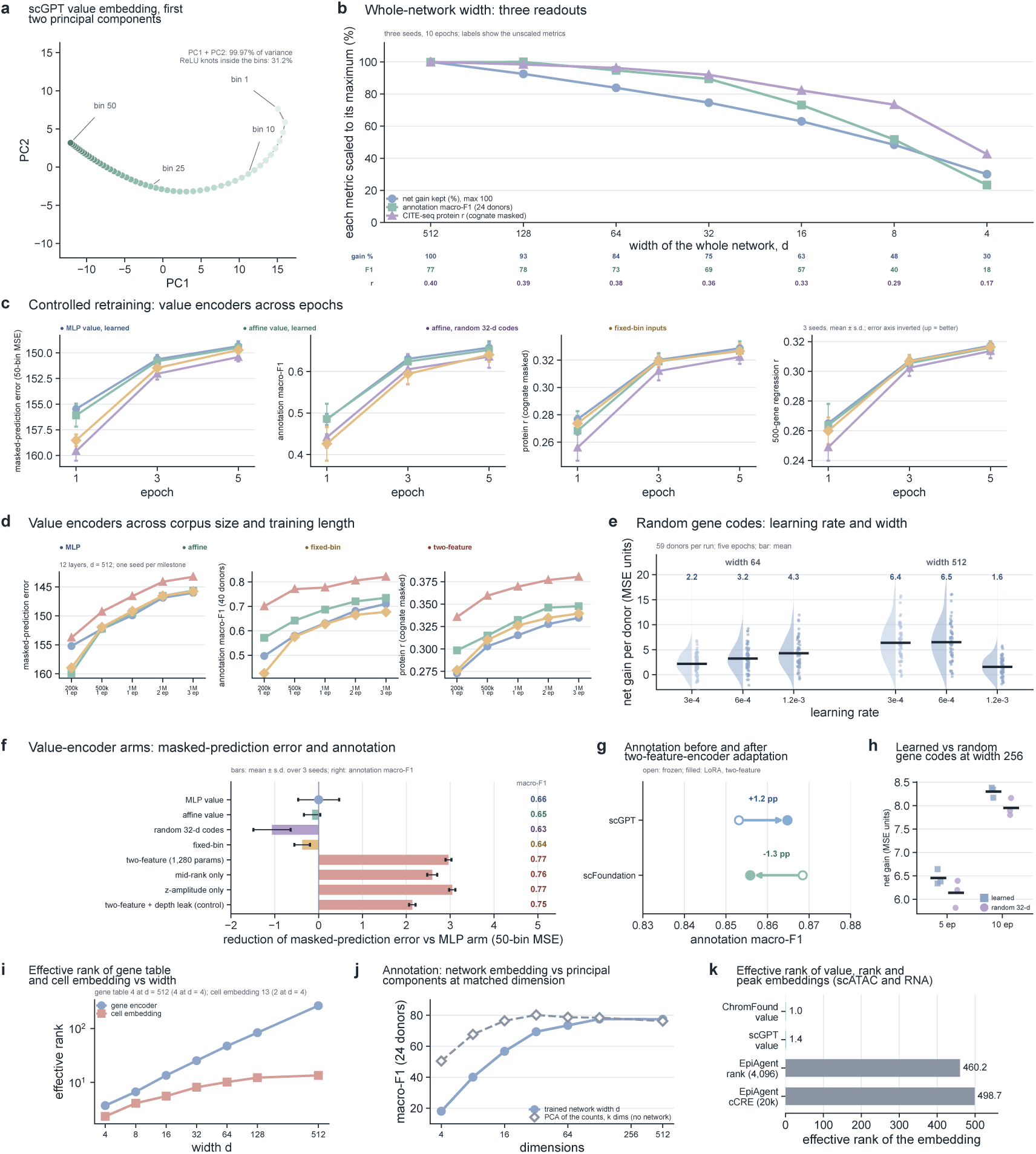
Value-encoder comparisons separate scalar encoding from backbone capacity. **a**, Pretrained scGPT value embeddings are projected onto principal components. Colour identifies input bins; annotations summarise geometry. **b**, Width scans compare masked prediction, cell-type annotation and surface-protein prediction. Curves are maximum-scaled; labels retain original units. **c**, Controlled Transformers compare value encoders during training. Bars show mean *±* standard deviation across three runs; error decreases upwards. **d**, Encoder comparisons span corpus scales and training durations. Symbols distinguish encoders. **e**, Random-code models compare learning rates and widths. Dots represent 59 donors per run; ticks mark means. **f**, Encoder ablations compare masked-prediction error and annotation. Bars show error reductions *±* standard deviation across three runs; labels give macro-F1. **g**, Pretrained scGPT and scFoundation compare cell-type annotation before and after two-feature-encoder adaptation. Open and filled circles distinguish states; arrows pair them. **h**, Learned and random gene codes compare net gain at matched width. Dots represent runs; ticks mark means. **i**, Gene-table and cell-embedding effective ranks are compared across widths. Bands show mean *±* standard deviation across three runs. **j**, Network embeddings and principal-component projections compare annotation at matched dimension. Line styles distinguish representations across 24 donors. **k**, Accessibility and RNA embeddings compare effective rank. Bars distinguish value, rank and peak-identity embeddings (Methods).

**Extended Data Figure 3.**
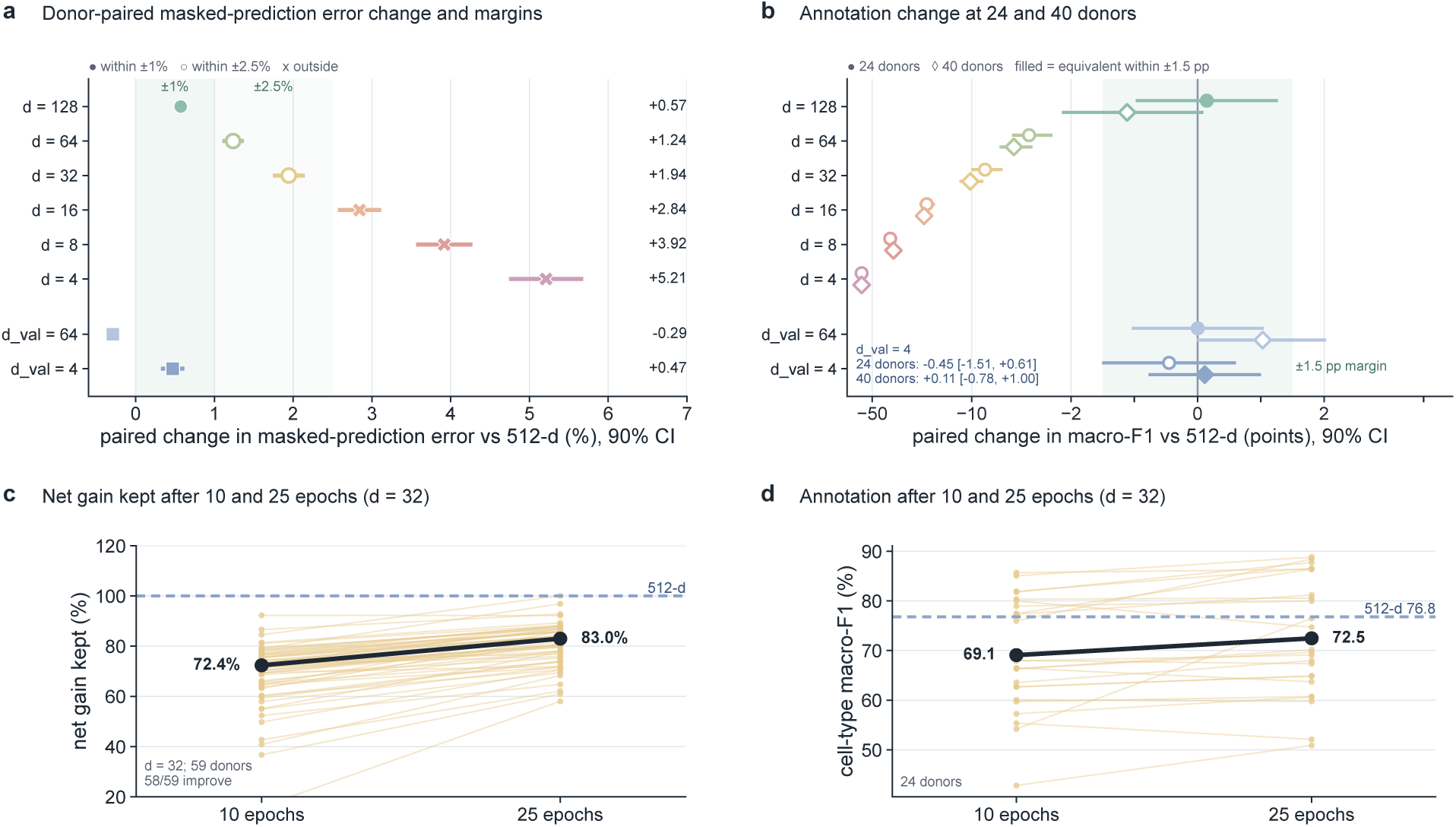
Equivalence tests distinguish value-track narrowing from whole-network narrowing. **a**, Donor-paired masked-prediction errors compare narrowed models with their wide references across 59 held-out donors. Bars show two-sided 90% paired-donor t intervals; circles denote whole networks and squares value tracks, with filled markers passing *±*1%, open markers passing *±*2.5%, and crosses failing both margins. **b**, Cell-type annotation contrasts compare two donor protocols. Circles represent 24 donors and diamonds 40 donors; bars show two-sided 90% paired-donor t intervals, shading marks *±*1.5 points and filled symbols denote equivalence within that margin. **c**, Longer training is evaluated for a narrow whole network on masked prediction. Thin lines pair donors, the thick line joins pooled net gains, and the dashed line marks the wide reference. **d**, The same training comparison evaluates cell-type annotation across 24 held-out donors. Thin lines pair donor scores and the thick line joins means; the dashed line marks the wide model (Methods).

**Extended Data Figure 4.**
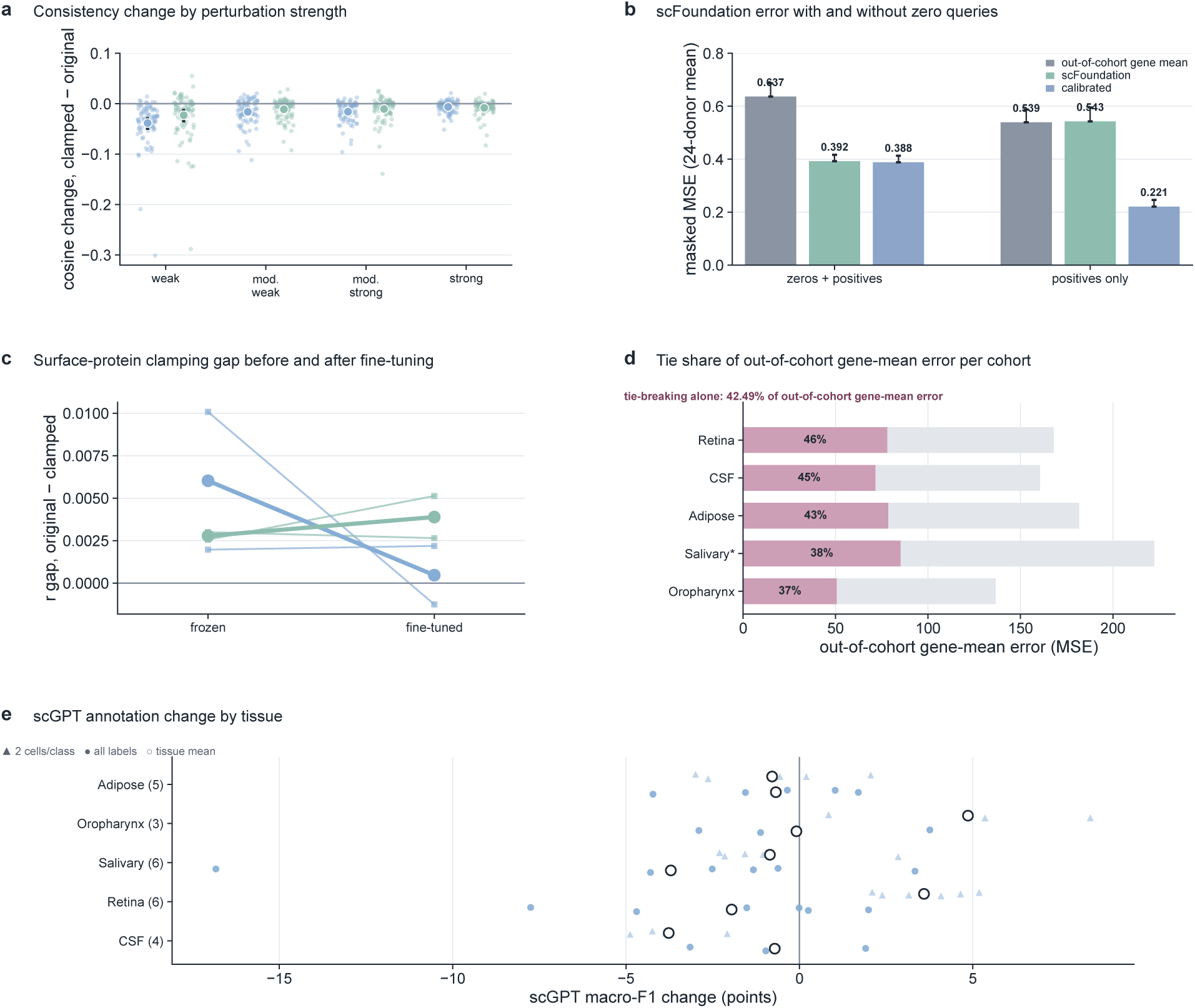
Clamping effects depend on the readout and evaluation population. **a**, Perturbation-response consistency compares clamping with original input across perturbation-strength quartiles. Dots represent individual perturbations among 320 targets; summary points show mean cosine changes with 95% perturbation-bootstrap intervals. **b**, scFoundation masked prediction compares mixed zero–positive and positive-only query protocols across 24 donors. Bars distinguish gene means, raw predictions and calibrated predictions; upward error bars show standard errors of donor means. **c**, Surface-protein prediction compares original-minus-clamped correlation gaps before and after partial fine-tuning. Thin lines connect cohort gaps across four held-out donors; thick lines connect cohort means, with colour identifying models. **d**, Rank-bin gene-mean error is partitioned by cohort. Coloured bar segments show analytical tie variance within total error; the remaining segments show error beyond ties. **e**, Pretrained scGPT cell-type annotation contrasts are grouped by tissue. Triangles mark two labelled cells per class, circles all labels, and rings tissue means; labels give donor counts (Methods).

**Extended Data Figure 5.**
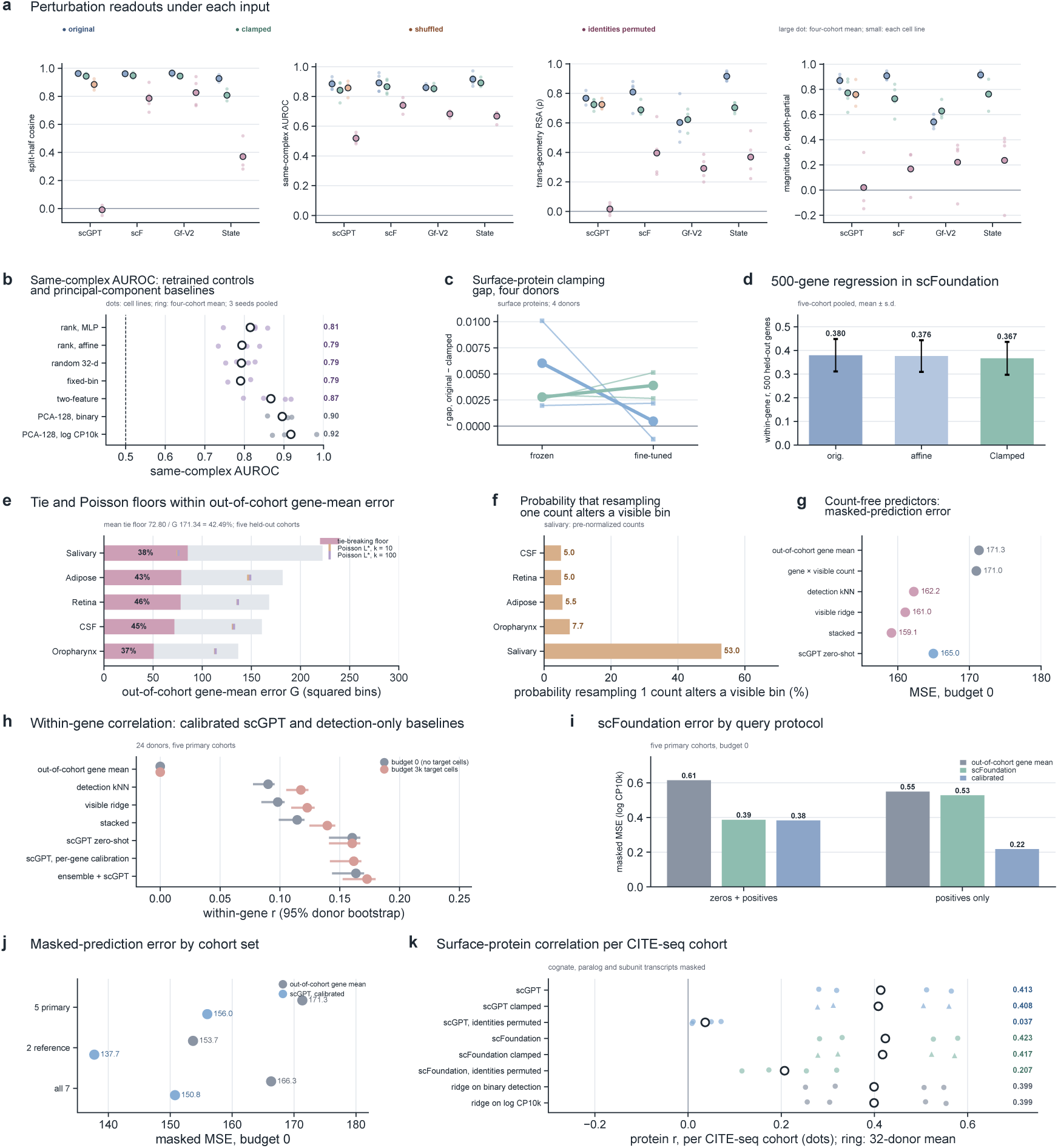
Count noise and calibration shape masked-prediction and transfer readouts. **a**, Perturbation consistency, complex recovery, response geometry and depth-adjusted magnitude compare input interventions. Small dots denote cell lines; large dots denote four-cohort means. **b**, Controlled and principal-component baselines compare protein-complex recovery. Dots mark cell lines; rings mark mean area under the receiver operating characteristic curve (AUROC). **c**, Fine-tuning compares surface-protein clamping gaps across four donors. Thin lines denote cohorts; thick lines denote means. **d**, scFoundation input variants compare held-out-gene prediction. Bars show five-cohort mean within-gene correlation *±* standard deviation. **e**, Gene-mean error is compared with noise floors. Bars show total and tie error; ticks mark Poisson floors under alternative neighbourhood sizes. **f**, Single-query count resampling tests visible-bin stability. Bars show cohort probabilities; salivary counts are pre-normalised. **g**, Count-free predictors compare masked-prediction error with scGPT. Points show aggregate errors without target-cohort fitting. **h**, Pretrained scGPT calibration compares within-gene correlation across fitting budgets. Points show estimates; bars show 95% stratified donor-bootstrap intervals. **i**, scFoundation calibration compares query protocols. Bars show error in log counts per ten thousand (CP10k). **j**, Temporal cohort subsets compare masked prediction. Colour distinguishes gene means from calibrated scGPT. **k**, Surface-protein prediction compares model inputs and count baselines. Dots denote cohorts; rings show 32-donor means with cognate-related transcripts masked (Methods).

**Extended Data Figure 6.**
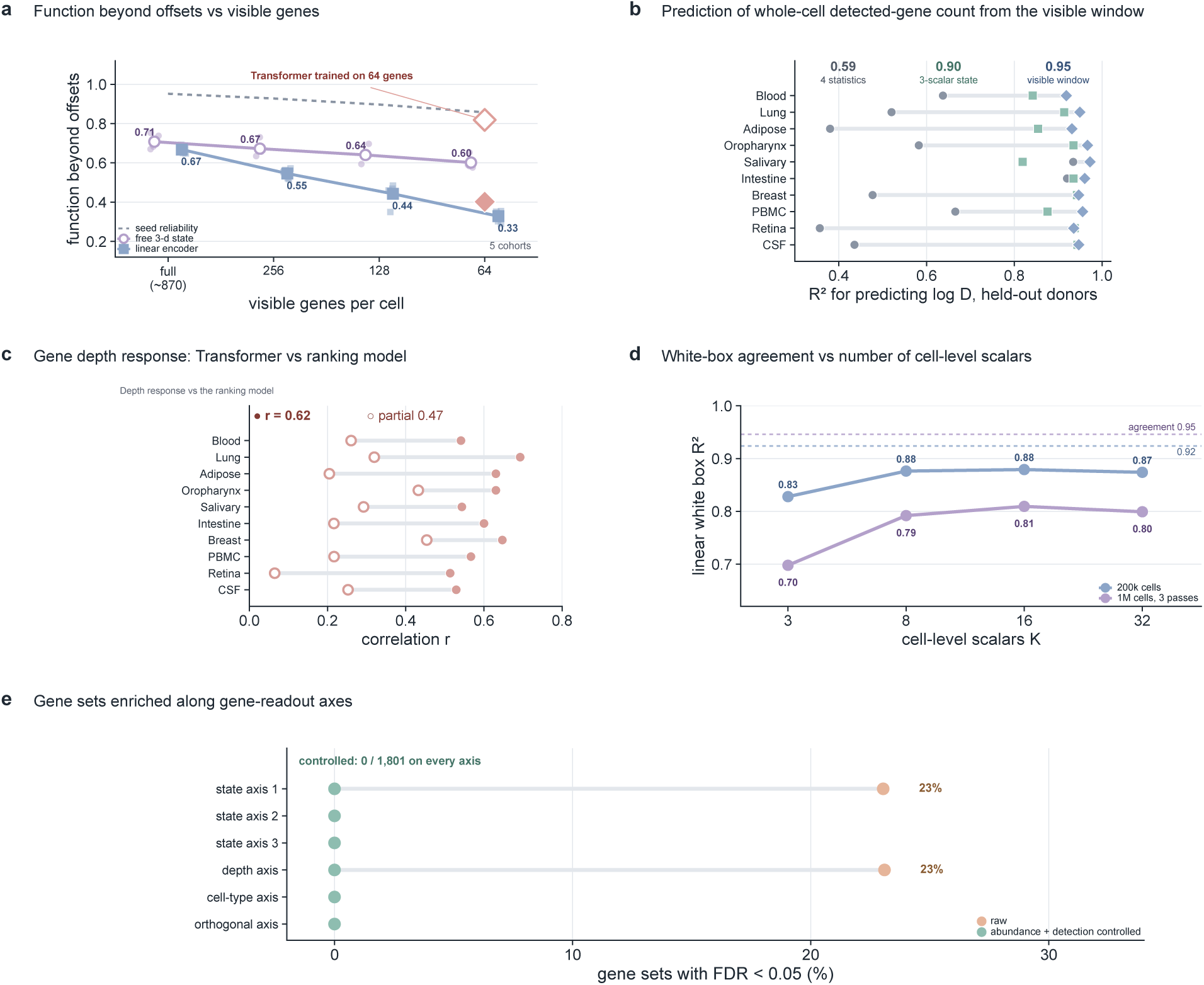
Sparse context limits linear recovery of a low-dimensional cell state. **a**, White-box fits compare progressively subsampled visible contexts across five cohorts. Small points denote cohorts; curves distinguish linear encoders, free states and independent-model reliability, while diamonds mark models trained with sparse context. **b**, Visible-input features predict whole-cell detected-gene count on held-out donors. Symbols distinguish window features, cell state and summary statistics across ten cohorts; annotations give pooled agreement. **c**, Gene-specific depth responses are compared with a multinomial ranking model. Filled and open points distinguish raw correlations from correlations adjusted for abundance and detection; rows identify cohorts. **d**, Linear white-box agreement is compared across numbers of cell-level scalars and training scales. Lines identify training conditions; dashed horizontal references mark independent-model agreement. **e**, Gene-readout axes are tested for gene-set enrichment before and after abundance control. Paired points show significant-set shares among 1,801 sets, using a false-discovery-rate threshold of 0.05 (Methods).

**Extended Data Figure 7.**
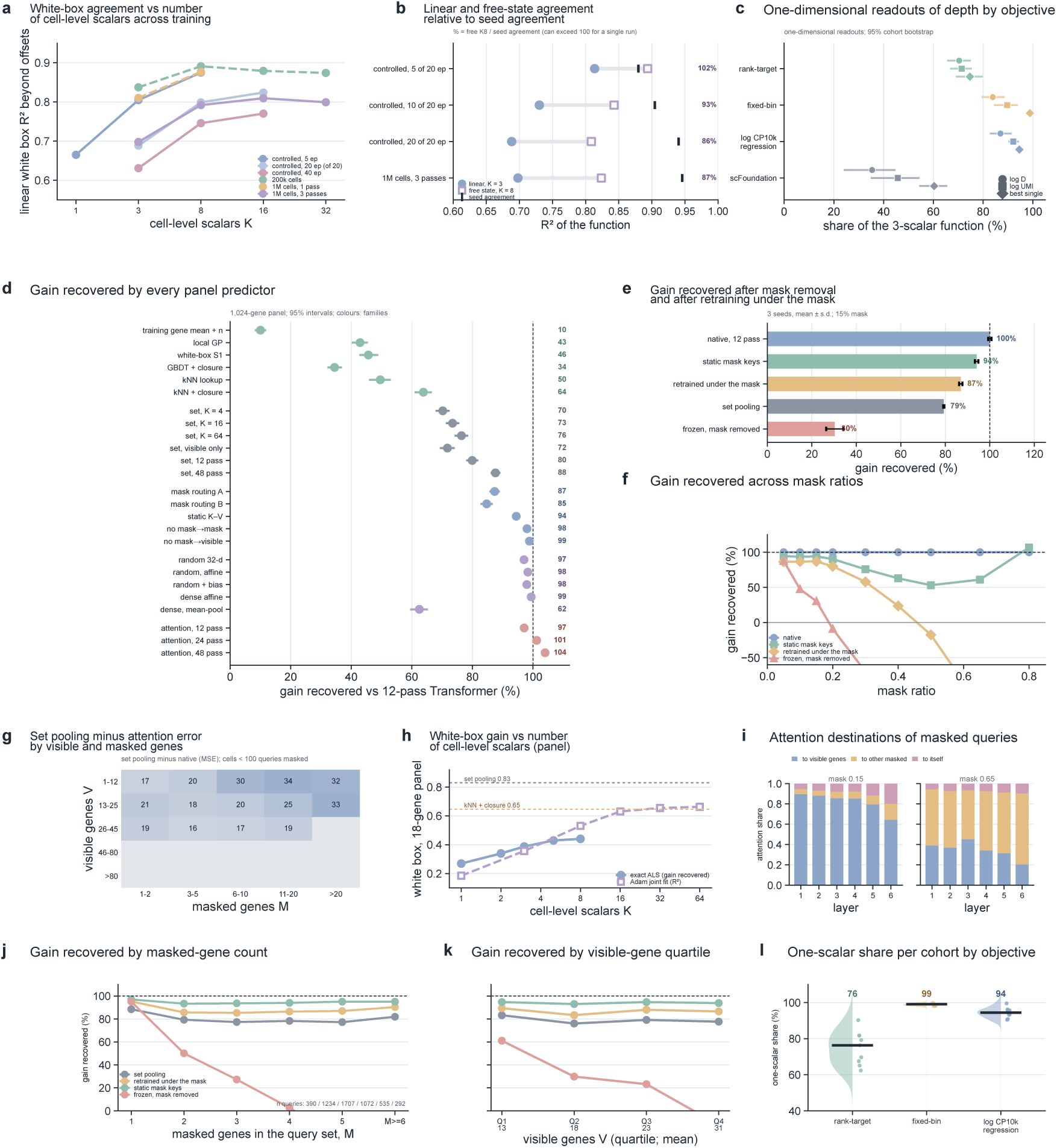
Sparse-panel controls distinguish attention routing from state dimensionality. **a**, White-box dimensionality curves compare training conditions. Lines identify schedules and corpus sizes. **b**, Linear and free-state fits compare model agreement. Symbols distinguish fits; ticks mark independent-run agreement. **c**, Depth covariates compare objectives. Symbols identify covariates; bars show 95% cohort-bootstrap intervals. **d**, Sparse-panel predictors compare masked-prediction gain. Colours identify families; bars show 95% run-by-cell bootstrap intervals. **e**, Attention masks compare post-hoc removal with retraining. Bars show mean *±* standard deviation across three runs. **f**, Mask-ratio sweeps compare routing variants. Curves show gain relative to the reference Transformer. **g**, Attention advantage is grouped by visible and masked-gene counts. Colour shows error differences; grey excludes bins below 100 queries. **h**, White-box fits compare numbers of cell-level scalars. Solid curves show alternating-least-squares gain share; dashed curves show joint-fit distillation agreement. **i**, Masked-query attention is partitioned by destination. Stacks distinguish visible genes, other queries and self-attention at each layer. **j**, Routing variants compare gain across masked-query-count strata. Curves distinguish variants; labels give query counts. **k**, The same variants compare visible-context quartiles. Tick labels give quartiles and mean visible-gene counts. **l**, Single-scalar contributions compare objectives across nine cohorts. Dots denote cohorts; ticks mark medians (Methods).

**Extended Data Figure 8.**
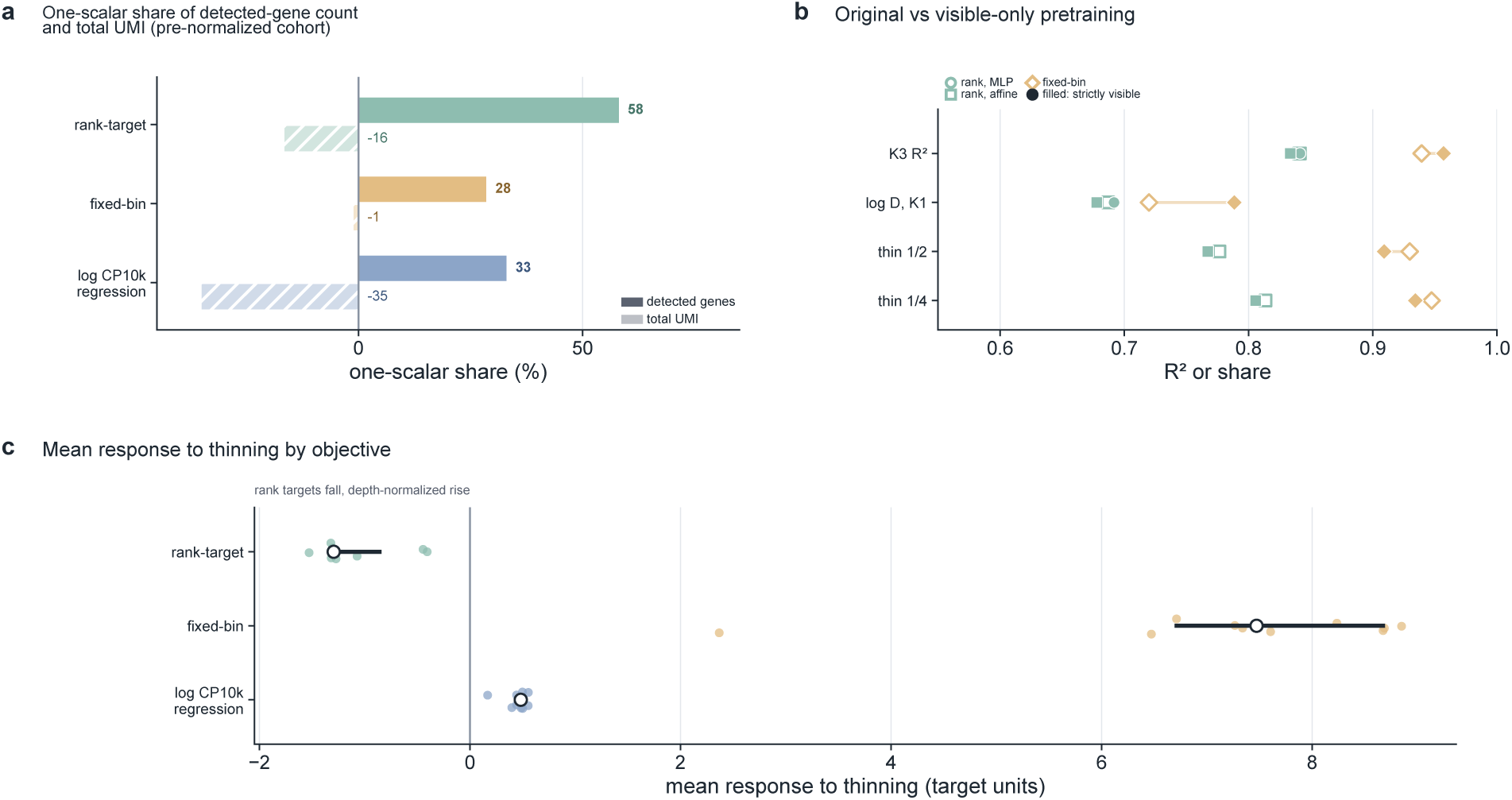
Objective and input construction determine which depth covariate describes the response. **a**, Depth predictors compare objectives in the pre-normalised salivary-gland cohort. Solid bars show detected-gene-count contributions and pale hatched bars total unique molecular identifier (UMI) contributions, each expressed relative to the cell-state increment beyond gene offsets. **b**, Original and visible-only pretraining compare white-box fits and thinning responses. Open and filled markers distinguish input construction, connectors pair conditions, and marker shape distinguishes rank-target encoders from fixed-bin log counts per ten thousand (CP10k) encoders; rows report white-box agreement, detected-gene-count contribution and response agreement at both thinning doses (Methods). **c**, Mean thinning responses compare rank-target, fixed-bin log CP10k and continuous-regression objectives. Coloured dots represent ten cohorts; rings mark cohort medians with 95% cohort-bootstrap intervals, and the vertical line marks zero response in each objective’s target units, with half- and quarter-depth interventions defined in the schematic (Methods).

**Extended Data Figure 9.**
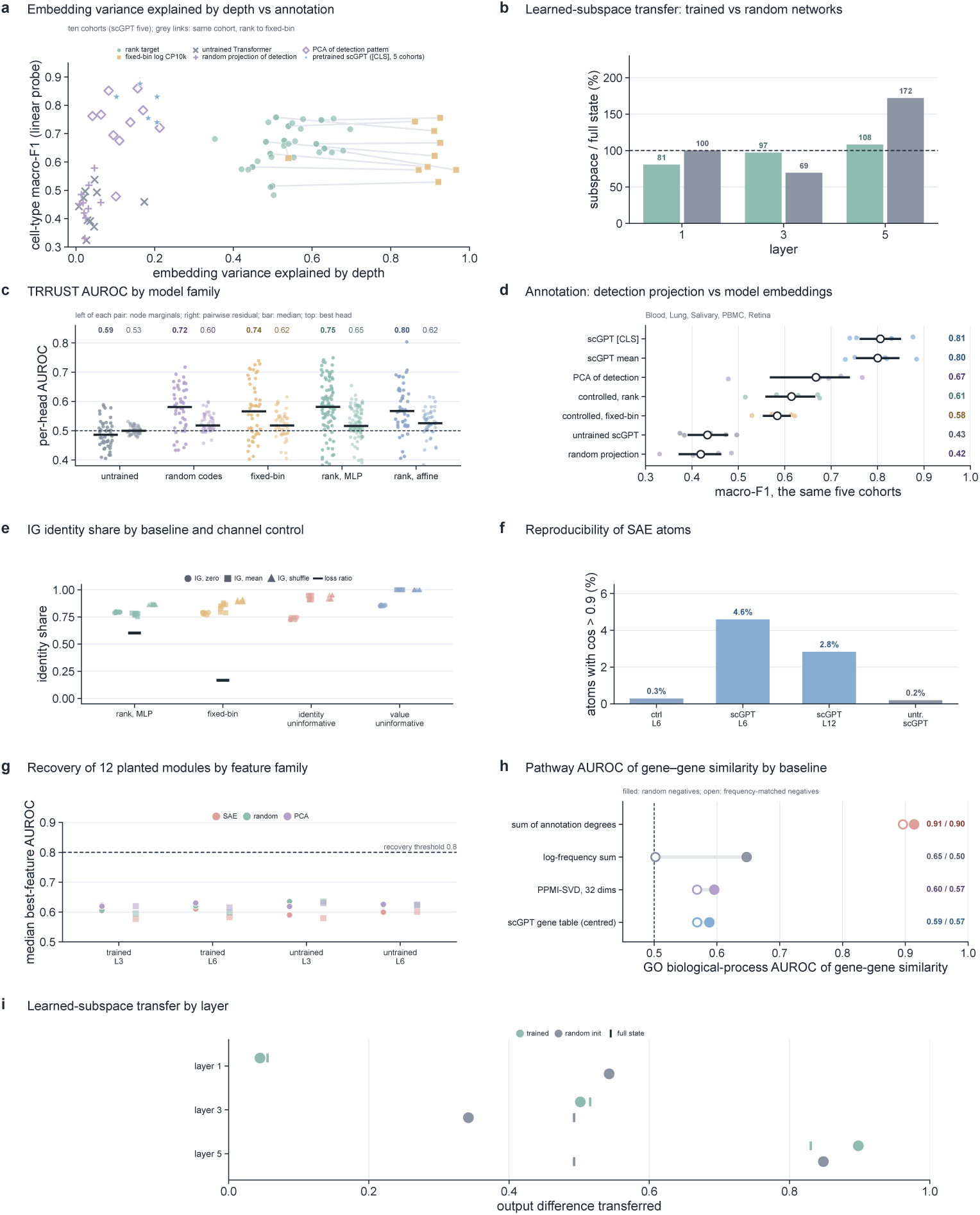
Matched nulls explain much of the structure recovered by interpretability readouts. **a**, Cell embeddings compare depth-explained variance with cell-type annotation. Symbols identify models across ten cohorts, five for pretrained scGPT; links pair objectives. **b**, Learned-subspace swaps compare trained and randomly initialised networks. Bars show transfer relative to each network’s full-state swap. **c**, Attention components compare regulatory-interaction recovery. Dots mark heads; ticks mark median area under the receiver operating characteristic curve (AUROC), pairing marginals with residuals. **d**, Detection projections and model embeddings compare annotation across five cohorts. Rings show means with 95% cohort-bootstrap intervals. **e**, Integrated Gradients (IG) compares informative and uninformative channels. Shapes identify attribution baselines; black ticks mark defined intervention-loss identity shares. Zero- or float32-resolution-denominator ratios are omitted while their IG points are retained; the ratio is not a causal truth for IG. **f**, Sparse autoencoder (SAE) atoms compare reproducibility across fits. Bars show shares exceeding cosine similarity 0.9. **g**, Feature families compare recovery of twelve planted modules. Shapes distinguish value conditions, colours feature families, and the dashed line the AUROC 0.8 recovery threshold. **h**, Gene-similarity baselines compare pathway recovery. Filled and open markers distinguish random and frequency-matched negatives. **i**, Learned-subspace transfer compares trained and randomly initialised networks across layers. Dots show subspace transfer; ticks show full-state references (Methods).

**Extended Data Figure 10.**
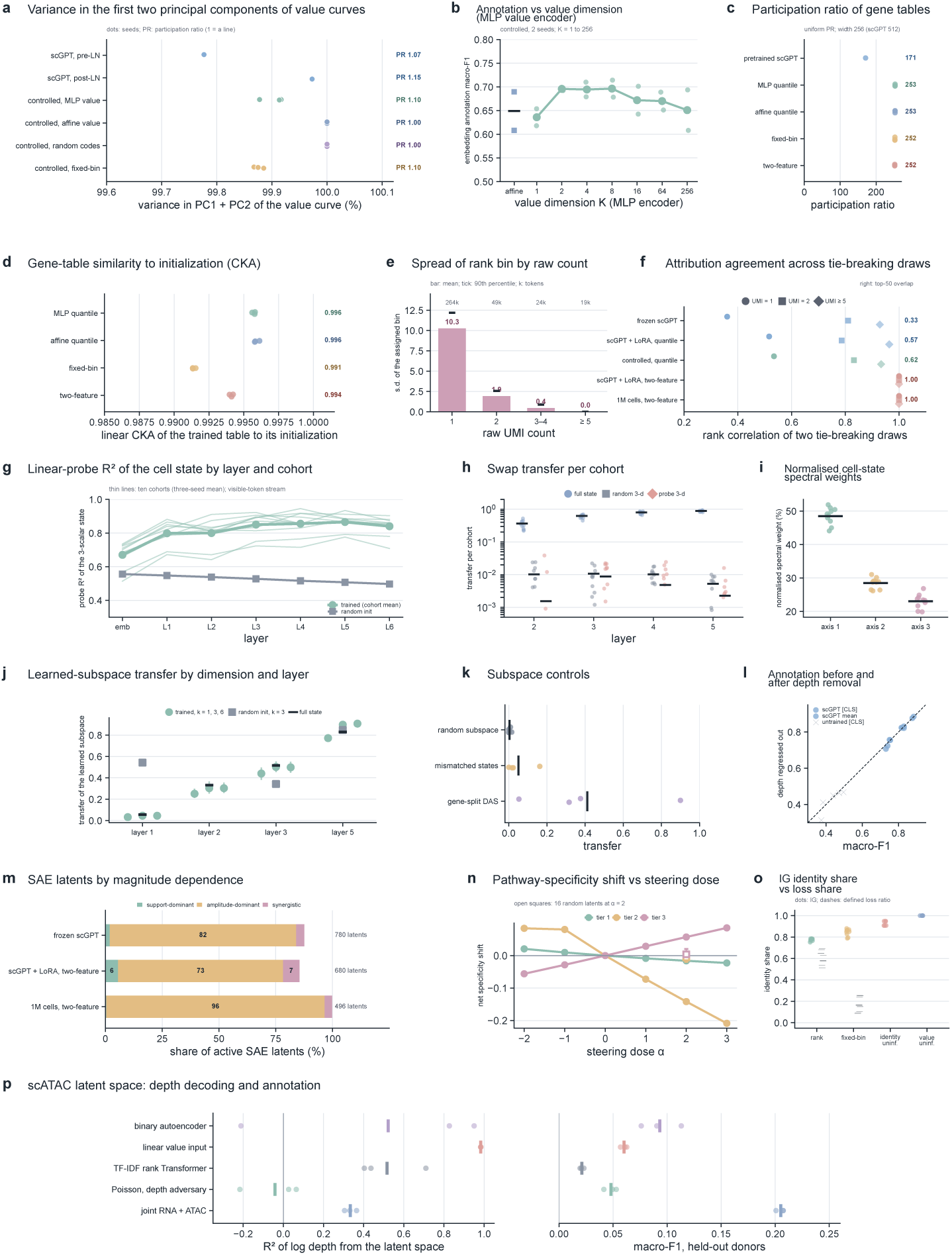
Geometry and intervention controls distinguish readable structure from functional use. **a**, Value curves compare geometry. Dots show variance shares and participation ratios. **b**, Value dimensions compare cell-type annotation. Dots mark two runs. **c**, Gene tables compare effective rank. Dots show participation ratios. **d**, Gene tables compare initialisation similarity. Dots show centred kernel alignment. **e**, Unique molecular identifier (UMI) strata compare bin dispersion. Bars mark means; ticks mark ninetieth percentiles. **f**, Gradient-times-input attribution compares tie-breaking draws. Shapes identify UMI strata; labels give top-50 overlap. **g**, Cell-state probes compare layers. Lines distinguish cohorts and model summaries. **h**, Swap interventions compare transfer. Log-scaled dots mark cohorts; ticks mark pooled slopes. **i**, Cell-state spectra compare cohorts. Dots show normalised spectral weights. **j**, Distributed alignment search compares transfer. Bars show 95% bootstrap intervals. **k**, Subspace controls compare transfer. Dots show conditions; ticks show means. **l**, Depth removal compares annotation. Dots compare cohort scores against equality. **m**, Sparse autoencoder (SAE) latents compare magnitude dependence. Stacks identify support-dominant, amplitude-dominant and synergistic tiers. **n**, Latent steering compares pathway specificity. Squares show means *±* standard deviation for 16 random latents at dose two. **o**, Integrated Gradients (IG) compares identity shares. Dots mark IG; ticks mark defined intervention-loss shares. Undefined ratios are omitted while IG points remain; the two measures use different endpoints. **p**, Chromatin-accessibility models compare depth decoding and annotation. Dots mark three runs; ticks mark means (Methods).

